# Global patterns and rates of habitat transitions across the eukaryotic tree of life

**DOI:** 10.1101/2021.11.01.466765

**Authors:** Mahwash Jamy, Charlie Biwer, Daniel Vaulot, Aleix Obiol, Homgmei Jing, Sari Peura, Ramon Massana, Fabien Burki

## Abstract

The successful colonisation of new habitats has played a fundamental role during the evolution of life. Salinity is one of the strongest barriers for organisms to cross, which has resulted in the evolution of distinct marine and terrestrial (including both freshwater and soil) communities. Although microbes represent by far the vast majority of eukaryote diversity, the role of the salt barrier in shaping the diversity across the eukaryotic tree is poorly known. Traditional views suggest rare and ancient marine-terrestrial transitions, but this view is being challenged by the discovery of several recently transitioned lineages. Here, we investigate habitat evolution across the tree of eukaryotes using a unique set of taxon-rich environmental phylogenies inferred from a combination of long-read and short-read metabarcoding data spanning the ribosomal DNA operon. Our results show that overall marine and terrestrial microbial communities are phylogenetically distinct, but transitions have occurred in both directions in almost all major eukaryotic lineages, with at least 350 transition events detected. Some groups have experienced relatively high rates of transitions, most notably fungi for which crossing the salt barrier has most likely been an important aspect of their successful diversification. At the deepest phylogenetic levels, ancestral habitat reconstruction analyses suggest that eukaryotes may have first evolved in non-saline habitats, and that the two largest known eukaryotic assemblages (TSAR and Amorphea) arose in different habitats. Overall, our findings indicate that crossing the salt barrier has played an important role in eukaryotic evolution by providing new ecological niches to fill.

## Main text

Adapting to new environments with very different physicochemical properties represent large evolutionary steps. When successful, habitat transitions can be important drivers of evolution and trigger radiations^1–4^. The marine-terrestrial boundary (here terrestrial encompassing both freshwater and soil^5,6^)—the so-called salt barrier—is considered one of the most difficult barriers to cross, because salinity preference is a complex trait that requires the evolution of multi-gene pathways for physiological adaptations^7–10^. These adaptations have been best studied in macroorganisms, for which the recorded marine-terrestrial transitions are few^11–13^. Microbes (prokaryotic and eukaryotic) are also typically regarded as infrequently crossing the salt barrier in spite of much larger population sizes and high dispersal ability^12,14^, but the role of the salt barrier as an evolutionary driver of microbial diversity remains poorly understood. For bacteria, higher habitat transition rates than anticipated have been reported^15^. For microbial eukaryotes, which represent the vast majority of eukaryotic diversity, no data exist to infer the global patterns and rates of habitat transitions at a broad phylogenetic scale. Extant marine and terrestrial eukaryotic communities are distinct in terms of composition and abundance of taxa^6,16^, a pattern that has been attributed to rare and ancient transitions between marine and terrestrial environments^14,17–22^. However, increasing inferences of recent transitions in specific clades such as dinoflagellates suggest that the strength of the salt barrier might not be as strong as previously envisioned^23–26^.

In this study, we used a unique hybrid approach combining high-throughput long-read and short-read environmental sequencing to infer habitat evolution across the eukaryotic tree of life. We newly generated over 10 million long environmental reads (ca. 4500 bp of the ribosomal DNA operon) from 21 samples spanning marine (including euphotic and aphotic ocean zones), freshwater, and soil habitats. The increased phylogenetic signal of long-reads allowed us to establish, together with a set of phylogenomics constraints, a broad evolutionary framework for the environmental diversity of eukaryotes. We then incorporated existing, massive short-read data (∼234 million reads) from a multitude of locations around the world to complement the taxonomic and habitat diversity of our dataset. With this combined dataset, we inferred the frequency, direction, and relative timing of marine-terrestrial transitions during the evolution of eukaryotes; we investigated which eukaryotic lineages are more adept at crossing the salt barrier; and finally, we reconstructed the most likely ancestral habitats throughout eukaryote evolution, from the root of the tree to the origin of all major eukaryotic lineages. Our analyses represent the most comprehensive attempt to leverage environmental sequencing to infer the evolutionary history of habitat transitions across eukaryotes.

## Results

### Long-read metabarcoding to obtain a comprehensive environmental phylogeny

A range of samples collected globally from marine and terrestrial habitats were deeply sequenced with PacBio (Sequel II) to obtain a comprehensive long-read metabarcoding dataset spanning the broad phylogenetic diversity of eukaryotes. These samples covered all major ecosystems, including the marine euphotic and aphotic zones (surface/deep chlorophyll maximum, and mesopelagic/bathypelagic, respectively), freshwater lakes and ponds as well as tropical and boreal forest soils (see Supplementary Table 1 for details). In total, we obtained 10.7 million Circular Consensus Sequence (CCS) reads spanning ∼4500 bp of the ribosomal DNA (rDNA) operon, from the 18S to the 28S rDNA genes. After processing, sequences were clustered into Operational Taxonomic Units (OTUs) within each sample at 97% similarity, resulting in 16,821 high-quality OTUs. To assess the potential biases of long-read amplicon sequencing, we performed a direct comparison with Illumina data (for the V4 and V9 hypervariable regions of the rDNA gene, and 18S reads extracted from metagenomic data) previously obtained for the same DNA from three marine samples^27^. This comparison revealed that our long-range PCR assay followed by PacBio sequencing retrieved similar eukaryotic community snapshots, with most groups detected at comparable abundances (Supplementary Figures 1-2). Additionally, the PacBio datasets detected several taxonomic groups that are absent from the V4 and V9 datasets. Importantly, over 80% of the V4 sequences were identical to the PacBio OTUs, indicating that our protocol for CCS processing generates high-fidelity data comparable to classical short-read metabarcoding (Supplementary Figure 1).

We then used a phylogeny-aware method to label all OTUs with appropriate taxonomic information^28^ (see Materials and Methods for details), and reconstructed a global eukaryotic phylogeny of environmental diversity based on the 18S-28S rDNA genes (Figure 1). In order to allow for transition rates to be estimated within a guiding taxonomic framework (see below), the major eukaryotic groups shown in Figure 1a were constrained to be monophyletic based on established relationships derived from phylogenomic inferences (reviewed in ^29^). These major lineages were defined as rank 4 in the taxonomic scheme of an in-house database derived from the protist ribosomal reference (PR2) database^30^ called *PR2-transitions*^31^. This phylogeny contains almost all known major eukaryotic lineages (Figure 1); most of the missing groups (e.g. kelp and seaweed) represent large multicellular organisms, or protists found in specific environments not sampled here (e.g. anoxic environments, see Supplementary Table 2). We also uncovered a proportion of novel diversity, i.e. OTUs highly dissimilar to reference sequences that are typically difficult to confidently assign to taxonomic groups. Long-read metabarcoding alleviates the issue of taxonomic assignment of highly diverging sequences, for example we found 863 sequences with <85% similarity to references in PR2 which were attributed a taxonomy based on their position in the tree, mostly belonging to apicomplexan parasites, fungi, and amoebozoans (Figure 1a and Supplementary Figure 3).

**Figure 1.**
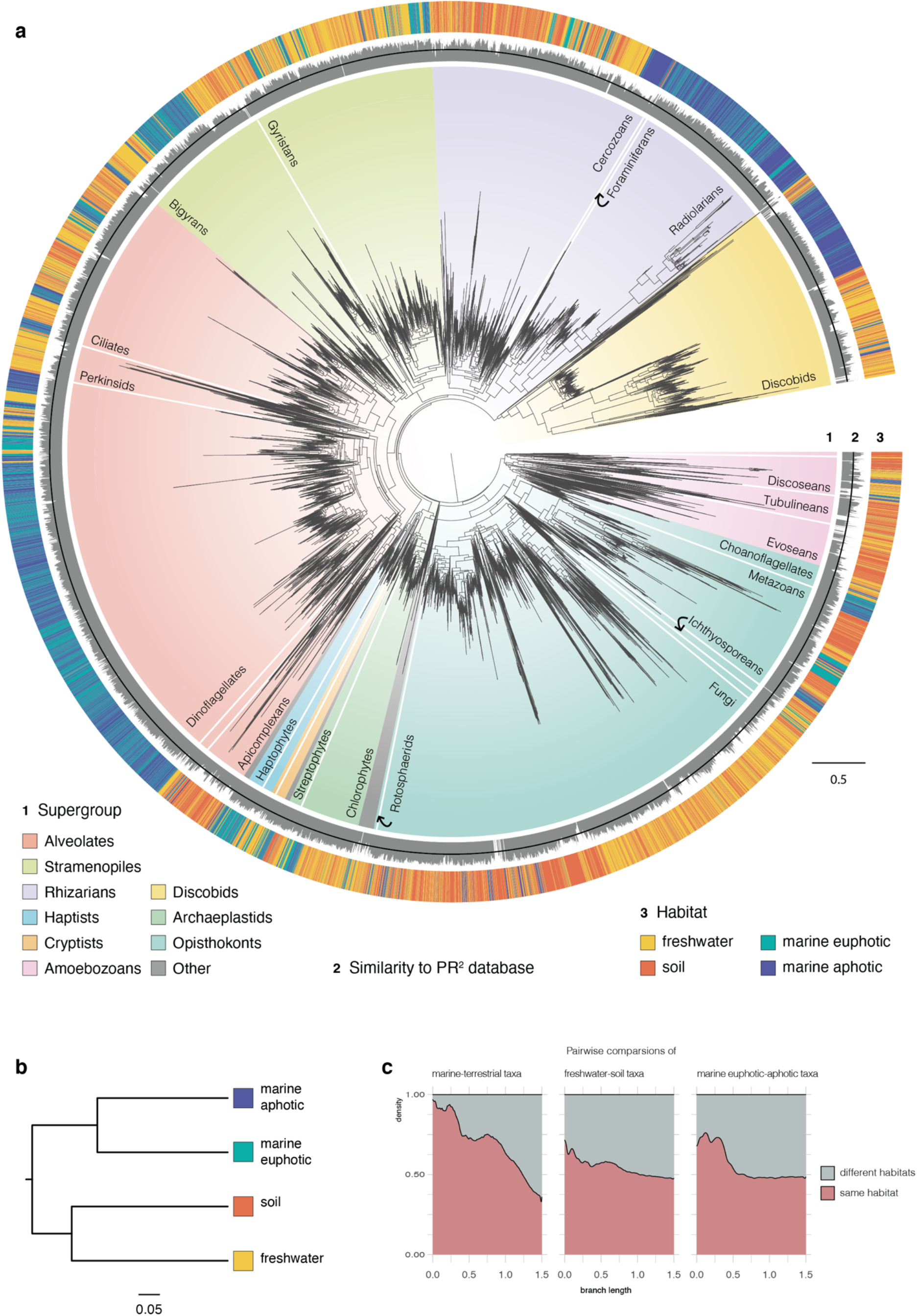
Global eukaryotic 18S-28S phylogeny from environmental samples and the distribution of habitats. **(a)** This tree corresponds to the best maximum-likelihood (ML) tree inferred using an alignment with 7,160 sites and the GTRCAT model in RAxML^32^. The tree contains 16,821 OTUs generated from PacBio sequencing of 21 environmental samples. The innermost ring around the tree indicates taxonomy, and the major eukaryotic lineages considered in this study are labelled. The second ring depicts percentage similarity with the references in the PR2 database and was set with a minimum of 70 and a maximum of 100, with the black line in the middle indicating 85% similarity. The third ring depicts which habitat each OTU belongs to. **(b)** Hierarchical clustering of the four habitats based on a phylogenetic distance matrix generated using the unweighted UniFrac method. (c) Stacked density plot of branch lengths between taxa pairs from the same or different habitats. Note that this plot should be interpreted with caution as each taxa-pair does not represent independent data-points due to phylogenetic relatedness.

### Detection of a salty divide in microbial eukaryotes

The global phylogeny in Figure 1 allows to visualize habitat preferences across the eukaryotic tree of life. Overall, we observed a clear phylogenetic distinction between marine and terrestrial lineages, with almost no OTU overlap between these two communities (Figure 1b-c; Unifrac distance = 0.959, p-value < 0.001). Within the marine and terrestrial biomes, soil and freshwater communities were found to be more distinct from each other (Unifrac distance = 0.76, p-value < 0.001) than the marine euphotic and aphotic communities (Unifrac distance = 0.64, p-value < 0.001) (Figure 1b and Supplementary Figure 4). However, we detected several sequences with high identity (>97% similar) present in the marine euphotic and aphotic samples (854 OTUs), and in the soil and freshwater samples (771 OTUs), suggesting that some taxa may be generalists in these sub-habitats (Figure 1c and Supplementary Figure 5).

We next sought to increase the number of samples and covered diversity by taking advantage of the mass of available short-read metabarcoding datasets. We gathered data from 22 studies conducted globally (including marine and terrestrial ecosystems), amounting to 234 million reads in total after processing (Supplementary Figure 6, Supplementary Table 3). We opted to use only the V4 region (ca. 260 bp) of the 18S rDNA gene as it was shown to have a greater phylogenetic signal than the V9 region^33^. The V4 reads were clustered into OTUs at 97% similarity for the marine euphotic (9977 OTUs), marine aphotic (2518 OTUs), freshwater (3788 OTUs), and soil (11935 OTUs) environments (Supplementary Table 4). These short-read OTUs were then phylogenetically placed onto the global long-read-based eukaryotic phylogeny using the Evolutionary Placement Algorithm (EPA)^34^ (Supplementary Figure 7), for which we compared the placement distributions for each sub-habitat. Interestingly, most placements occurred close to the tips of the reference tree, indicating that our long-read dataset adequately represents the diversity recovered by short-read metabarcoding (Supplementary Figure 7). Furthermore, the placement distributions for each habitat is consistent with our results based on the long-reads only, namely that marine and terrestrial communities are distinct, and at a finer level, soil and freshwater communities are more different from each other than communities in the surface and deep ocean (soil-freshwater earth mover’s distance = 1.14, marine euphotic-aphotic earth mover’s distance = 0.809; Supplementary Figure 8).

### Marine-terrestrial transition rates vary across major eukaryotic clades

The above results confirm that the salt barrier leads to phylogenetically distinct eukaryotic communities. We next asked how often have transitions between marine and terrestrial habitats occurred during evolution, which eukaryotic lineages have crossed this barrier more frequently, and in which direction? To answer these questions, we calculated habitat transition rates across the global eukaryotic phylogeny by performing Bayesian ancestral state reconstructions using continuous-time markov models^35^. We tested a null model, where transition rates from marine to terrestrial habitats (qMT) and vice versa (qTM) are constant throughout the eukaryotic phylogeny, against a heterogeneous model where qMT and qTM are estimated separately for each major eukaryotic lineage (illustrated in Figure 1). The null model had a posterior density of log-likelihoods with a mean of - 2008.45 (Supplementary Figure 9). Under this model, transitions from marine to terrestrial habitats are just as likely as the reverse across the tree. However, this general analysis hides important variations in habitat transition rates between groups, and indeed the heterogenous model presented a much better fit (log-likelihood score of -1819.91; Log Bayes Factor = 269.3; Supplementary Figure 9), indicating that habitat transition rates vary strongly across the tree.

To investigate in more detail the rate of habitat transition within each major eukaryotic group, we inferred taxon-rich clade-specific phylogenies by combining short-read data with the backbone phylogenies obtained from long-read data. Incorporating these short-read data allowed us to detect additional transition events that would have otherwise been missed with the long-read data alone (Supplementary Figure 10). We modelled habitat transition rates along clade-specific phylogenies containing both marine and terrestrial taxa that were sufficiently large (at least 50 tips) to get precise estimates. We also excluded discobid excavates and discosean amoebozoans as preliminary analyses showed ambiguous transition rate estimates owing to large phylogenetic uncertainty. Fungi were found to have by far the highest transition rates for a given amount of evolutionary change; we estimated around 90 expected transition events along a branch length of one substitution/site in the phylogeny. These results indicate that habitat shifts are associated with very little evolutionary change in the ribosomal DNA sequences (Figure 2a). After fungi, cryptophytes and gyristans (ochrophyte algae, oomycete parasites and several free-living flagellates) had the highest global rates (around 8.2 and 3.4 and expected transitions per substitution per site).

**Figure 2.**
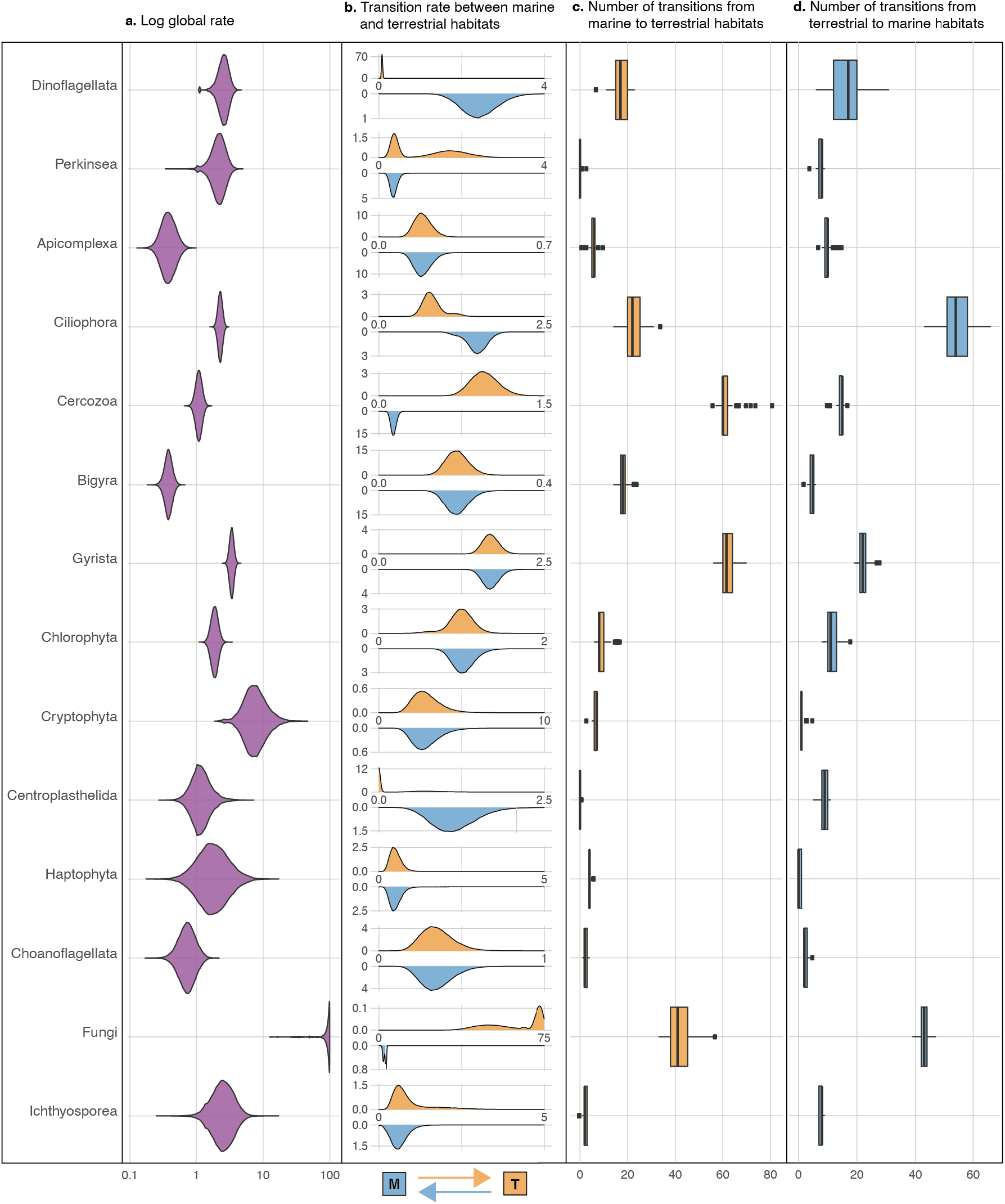
Habitat transition rates and number of transition events estimated for each major eukaryotic lineage. (a) Posterior probability distributions of the global rate of habitat evolution, which indicate the overall speed at which transitions between marine and terrestrial habitats have occurred in each clade regardless of direction. Rates were estimated along clade-specific phylogenies (see Supplementary Figure 10) using Markov Chain Monte Carlo (MCMC) in BayesTraits using a normalized transition matrix. (b) The posterior probability distribution of transition rates from marine to terrestrial habitats (top in orange), and from terrestrial to marine habitats (below in blue). (c) Number of transitions from marine to terrestrial habitats and (d) in the reverse direction for each clade as estimated by PASTML using Maximum Likelihood (see Materials and Methods for details).

At a finer phylogenetic resolution, several subclades within stramenopiles, as well as ciliates, seemed particularly adept at crossing the salt barrier, especially chrysophytes, diatoms, and spirotrich ciliates (11.8, 8.7, and 3.8 expected transition events per substitution per site respectively; Supplementary Figure 11-12). At the other extreme, groups such as bigyrans (heterotrophic stramenopiles related to gyristans) and apicomplexans (a group of parasites including the malaria pathogen) displayed the lowest habitat transition rates (around 0.4 expected transitions for every substitution per site). These results were further confirmed with sequence similarity network analyses, which showed high assortativity between marine and terrestrial sequences for bigyrans and apicomplexans (meaning that terrestrial and marine sequences formed distinct clusters at varying similarity thresholds), as opposed to gyristans and fungi, which showed low assortativity (Supplementary Figure 13).

Within each major eukaryotic group, we next inferred the frequency for each direction of the transitions between marine to terrestrial habitats. We found that all clades investigated had non-null transition rates in both directions, with the exception of centrohelids which had a terrestrial colonization rate that was not significantly different from zero in 99/100 trees used for calculation (Figure 2b). These results indicate that in nearly all major eukaryotic lineages containing terrestrial and marine taxa, transitions have occurred in both directions. Some clades had symmetrical transition rates, indicating that the tendency to colonize marine environments was not significantly different from the tendency to colonize terrestrial environments; this was for example the case of apicomplexans, bigyrans, gyristans, chlorophytes, cryptophytes, haptophytes, and choanoflagellates (Figure 2b). However, some groups showed marked differences in one direction or the other. Dinoflagellates, for example, show a much greater transition rate for colonizing marine habitats (about 31 times more likely). Ciliates have also transitioned more frequently towards marine environments, but the difference is smaller (1.8 times more likely). On the other hand, transitions to terrestrial environments were significantly more likely than the reverse direction for fungi and cercozoans (about 21.5 and 7.2 times more likely, respectively). Finally, the directionality of habitat transition appears to be heterogeneous also within the major eukaryotic groups (Supplementary Figures 14-17). Indeed, for some selected subclades such as ascomycetes and basidiomycetes within fungi, the transition rates to marine environments were higher as compared to non-Dikarya fungi, although fungi as a whole showed a marked tendency to colonize terrestrial habitats (qTM = 8.47 vs. 1.65 respectively; Supplementary Figure 17).

Finally, we estimated the number of transition events within each clade by generating discrete habitat histories using a maximum likelihood method^36^. We conservatively counted transition events only if they led to a clade with at least two taxa in the new habitat in order to distinguish between biologically active, speciating residents from wind-blown cells, resting spores or extracellular DNA from dead cells^37^. Our analyses revealed at least 350 transition events occurring over eukaryotic history, though the actual number is likely to be higher when considering lineages that have gone extinct. Out of these, 72 or more transition events occurred in fungi alone (39-47 transitions to marine environments detected, and 33-57 transitions to terrestrial environments detected) (Figure 2c-d). This was closely followed by gyristans and ciliates, with more than 60 putative switches each between environments (Figure 2c-d).

### Relative timing of habitat transitions during the evolution of the major eukaryotic groups

We next asked when during eukaryote evolution these transitions between marine and terrestrial habitats occurred. To calculate a relative timing for all marine-terrestrial transitions, we converted the clade-specific phylogenies into chronograms with relative dates (as in ^38^).

For each putative transition event, we measured the relative branch length from the inferred transition to the root of the clade. The general trend is that most transitions occurred relatively recently in the history of the groups (Figure 3). For instance, we detected no transition events in fungi older than 25% of the clade’s history, with the vast majority of all transitions occurring in the last 10% of the time that this group has been on earth. Assuming that fungi arose around 1 billion years ago^39–41^, this would imply that > 90% of all marine-terrestrial transitions (at least 63 transitions according to our analyses) in fungi occurred in the last 100 million years alone, with older transitions occurring predominantly towards marine environments. The observation that most transitions occurred towards present could be due to the increased challenges of inferring transition events early in the evolution of a group because of poorer resolution of deeper nodes due to little phylogenetic signal, and/or unsuccessful transitions leading to lineage extinctions in the new habitat. However, for a few clades at least (centrohelids, bigyra, apicomplexans, cercozoans, and chlorophytes), we detected a number of early transitions in the evolution of the group (Figure 3). Interestingly, the direction of these early habitat transitions is non-overlapping. For centrohelids and apicomplexans, the early transitions were mainly towards marine environments, possibly corresponding to repeated marine colonization events at the onset of the groups’ evolution. Early terrestrial colonization events were instead detected in cercozoans, chlorophytes, and bigyrans, together suggesting that early in the evolution of the major eukaryotic groups the pressure to move towards marine or terrestrial habitats was group-specific and directional.

**Figure 3.**
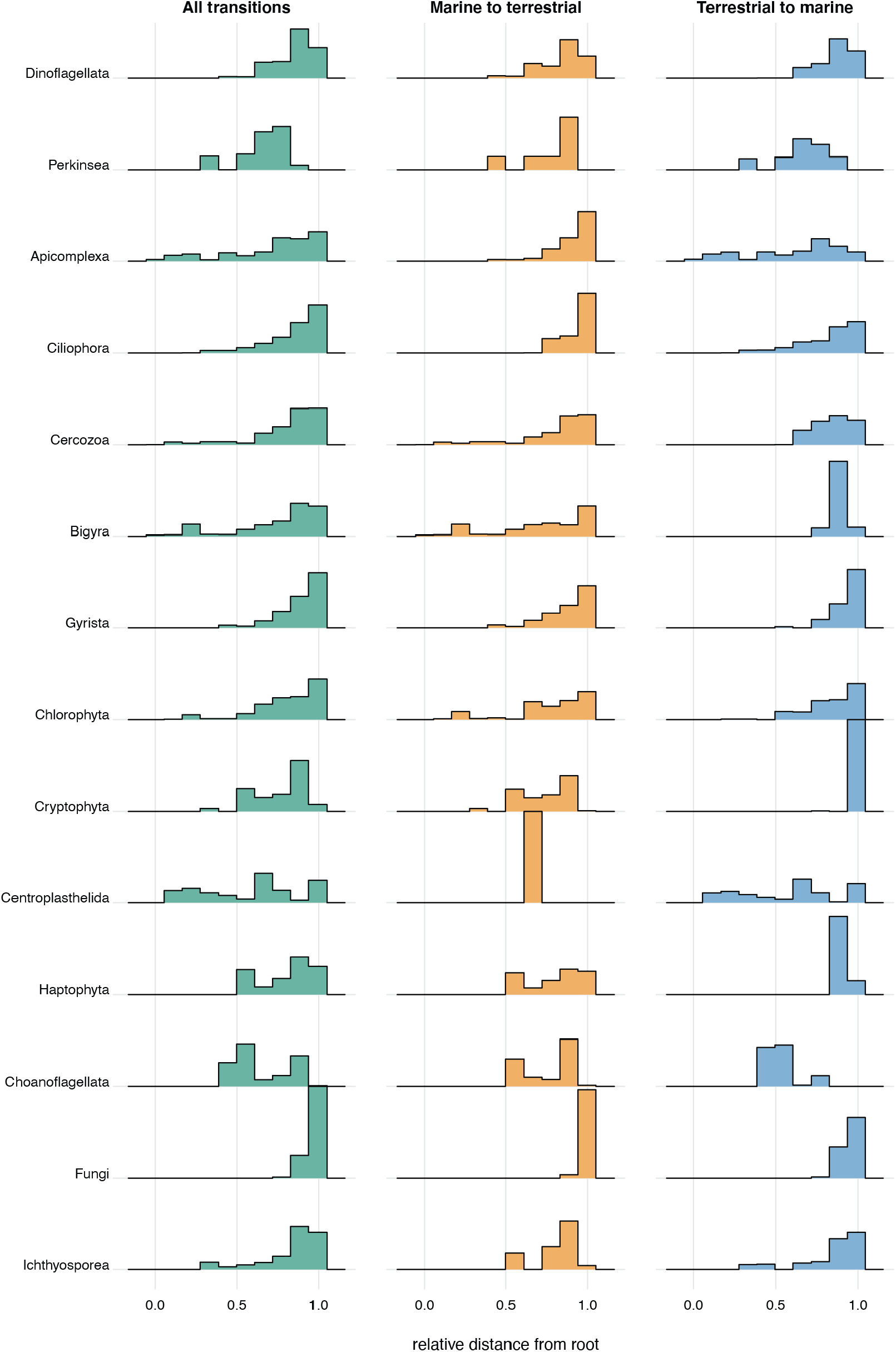
Ridgeline histogram plots displaying the timing of transition events as estimated from relative chronograms obtained with Pathd8^38^. The x-axis depicts the relative age for each clade.

### Ancestral habitat reconstruction of the major eukaryotic clades

Our global eukaryotic phylogeny of long-environmental OTUs, combined with the group-specific phylogenies including short-read metabarcoding data, represent a very dense set of environmental information put in a phylogenetic framework. We used this information to reconstruct in a Bayesian analysis the most likely ancestral environments from the root of the eukaryotic tree through the emergence of the major groups. Inferring the ancestral habitat of the last eukaryotic common ancestor (LECA) requires information about the root itself, which remains very contentious^29,42^. To accommodate uncertainties for the position of the root, we performed ancestral habitat reconstruction analyses using the two most commonly proposed root positions: (1) between the discobid excavates and all other eukaryotes^43^, and (2) between amorpheans (the group including animals, fungi and amoebozoans) and all other eukaryotes^44^. Both root alternatives converged towards the same habitats, suggesting with high confidence that LECA evolved in a terrestrial environment (Figure 4a).

**Figure 4.**
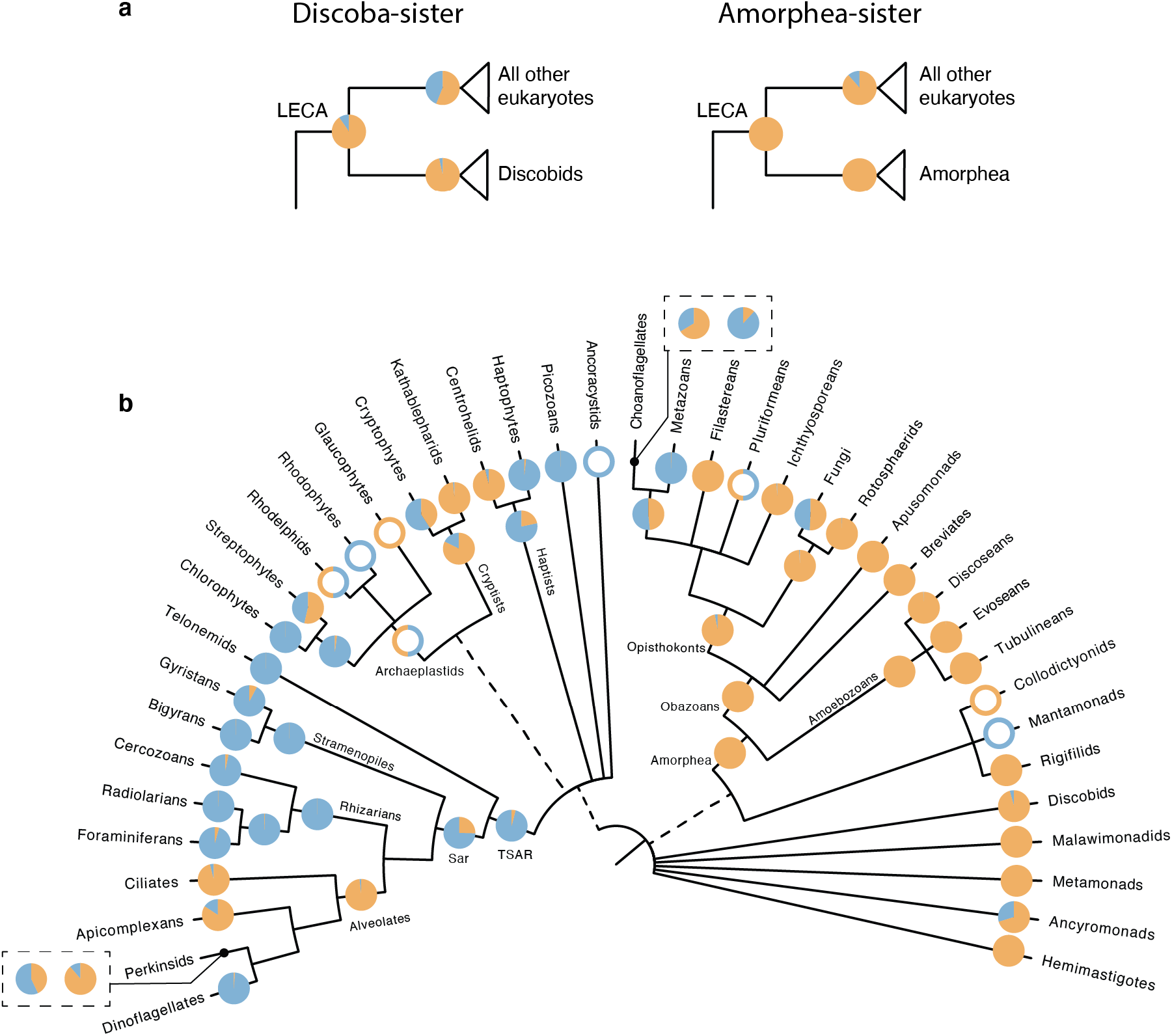
Ancestral states of major eukaryotic clades as estimated by BayesTraits on a set of 100 global PacBio phylogenies. Pie charts at each node indicate the posterior probabilities of likelihoods for the character states as follows: blue=marine, orange=terrestrial. Nodes with empty circles indicate wherever there was insufficient taxon sampling to infer ancestral habitats, but a reasonable estimate was made from existing literature (See Supplementary Note 1). (a) Ancestral habitat of the last eukaryotic common ancestor (LECA) as inferred using two different roots. (b) Ancestral states of major eukaryotic lineages. For the two cases where the incorporation of Illumina data inferred a different likely ancestral state, the results are shown in boxes. The pie chart on the right was obtained using the global eukaryotic phylogeny, while the pie chart on the left was obtained from clade-specific phylogenies. The tree is adapted from Burki et al^49^.

From this inferred terrestrial root, our analyses suggest that two of the largest mega-assemblages of eukaryotes, likely comprising more than half of all eukaryotic diversity^45^, arose in different environments. On one hand, the amorphean group likely originated in a terrestrial habitat (Figure 4b), where it initially diversified into obazoans (which include well-known lineages such as animals and fungi, but also several unicellular related lineages), as well as the amoebozoans. Consistent with previous studies, we inferred a marine origin for metazoans^46,47^, however for two obazoan lineages– fungi and the group containing metazoans and choanoflagellates–we could not determine a clear preference for their ancestral habitats. On the other hand, our analyses indicate that the expansive TSAR clade (containing the main eukaryotic phyla stramenopiles, alveoates, and rhiziarians, as well as the smaller group telonemids) most likely originated in a marine environment, following the transition of an ancestral population from a terrestrial root (Figure 4b). A marine origin is also likely for the major TSAR members, except for alveolates which were inferred to have a terrestrial origin.

Overall, the predicted ancestral habitats of most major eukaryotic clades match their current preferred habitat: this is for example the case for all amoebozoan lineages, radiolarians, dinoflagellates and foraminiferans. An exception is cercozoans for which a marine origin was inferred, but which now dominate terrestrial environments, particularly soils^6,48^. Interestingly, the results derived from the global eukaryotic phylogeny and the clade-specific phylogenies (which include short-read OTUs) were largely consistent except in two cases: the phylogeny of perkinsids changed the origin from terrestrial to marine for these parasites of animals, while the phylogeny of choanoflagellates switched from a marine to a terrestrial origin (Figure 4b).

## Discussion

In this study, we use a unique combination long- and short-read data to obtain an evolutionary framework of environmental diversity and infer habitat preference evolution across the eukaryotic tree. High-throughput long-amplicon sequencing followed by careful processing of the data provide high-quality sequences containing improved phylogenetic signal for the vast environmental diversity^28,50–52^. We generated over 10 million long-read metabarcoding data spanning the eukaryotic rDNA operon, which assembled into nearly 17,000 OTUs, for marine and terrestrial ecosystems. We then added two additional layers of phylogenetic information: (i) a much larger mass of available short-read metabarcoding data to more deeply cover the molecular diversity of environmental microbes, and (ii) a set of well-accepted constraints derived from published phylogenomic analyses to fix the backbone of our eukaryotic tree. By combining all this information, we show that we can infer evolutionary patterns at global scales across the tree.

We confirm that the salt barrier has been a major factor in shaping eukaryotic diversity^6,14,16^, and that marine-terrestrial transitions are infrequent in comparison to transitions across other habitats such as between freshwater and soil (Figure 1). Our analyses detected at least 350 transition events (Figure 2), although this number is likely to be higher when considering more ancient transitions that were not detected, and more recent transitions that will only be revealed by sequencing more locations (for e.g. in ^24^). These difficult-to-achieve environmental crossings have likely played important evolutionary roles by allowing colonizers to reach vacant ecological niches. For example, crossing the salt barrier may have led to the establishment of some major eukaryotic assemblages such as TSAR, or highly diverse lineages such as the oomycetes and vampyrellids (Supplementary Figure 12). Marine-terrestrial transitions have also allowed lineages such as diatoms, golden algae and spirotrich ciliates to expand their range to both habitats, contributing to the diversification of the vast eukaryotic diversity we see today. We unexpectedly found that 56% of the detected transitions occurred recently, in the last 10% of the evolutionary history of the respective groups (Figure 3), which is in contrast to a common idea that most marine-terrestrial transitions are ancient^14^. It is however unclear why colonization across the salt barrier would be more frequent in recent geological time, so this observation could instead be due to recent colonizing lineages having had less time to go extinct, and thus more likely to be represented in our generated phylogenies^53^. At its deepest phylogenetic level, our analyses suggest that the earliest eukaryotes inhabited terrestrial habitats (Figure 4) and not marine habitats as often assumed (e.g.^54–56^). While the fossil record for early eukaryotes is sparse and difficult to distinguish from prokaryotes, there is evidence for early eukaryotes in non-marine or low salinity environments from at least 1 Gyr ago^55^. Furthermore, other key early eukaryotic innovations, such as the origin of the plastid organelles, have been inferred to have occurred around 2 Gyr ago in low-salinity habitats^57,58^. Terrestrial environments are known to be more heterogenous^59^, and may thus have provided a wide range of ecological niches for early eukaryotes to occupy.

Our detailed investigation across the main groups of eukaryotes showed marked differences in the rates of crossing the salt barrier (200-fold globally). While some groups have low global transition rates, others show a higher tendency to cross this physiological barrier. Most notably, we inferred based on both the highest transition rates in our analysis and relatively high number of transition events (Figure 2), that fungi are the strongest eukaryotic colonizers between marine and terrestrial environments. This is consistent with previous studies documenting a multitude of close evolutionary associations between marine and terrestrial fungal lineages ^60–62^, which in turn suggests that many fungal species may be generalists that can tolerate a wide range of salinities^63,64^. Interestingly, fungi showed a much greater trend (21-fold) for colonizing terrestrial environments, where they are dominant, than the reverse. Whether this reflects a strong preference for terrestrial environments, or instead unequal diversification rates in the two habitats^65,66^, or both, is unclear and should be further investigated.

The differences in habitat transition rates across eukaryotes are likely the result of varying salinity tolerance that has prevented successful colonization events during the evolution in some groups. Among algae, comparative genomics showed large differences in gene content between marine and freshwater species, notably for ion transporters and other membrane proteins that likely play important roles in osmoregulation^67^. These different gene contents may be due, at least in part, to laterally acquired genes (LGT) that could facilitate successful crossing of the salt barrier, as proposed for other environmental adaptations^68–72^. In bacteria, it was hypothesized that particle-associated species can more easily cross the salt-barrier due to increased chances of acquiring osmoregulation-related genes through LGT^73^. Altogether, these observations raise the question of how protists in general acquire these genes, for example groups like diatoms (Supplementary Figures 11-12) which showed multiple transitions in both directions, and whether it is through LGT (as has been shown for some halophilic protists^68^), gene duplication, or through re-wiring of existing metabolic pathways (as shown for the SAR11 bacteria^74^). Other ecological factors also likely play a role as a colonizing organism does not only need to adapt to a different salinity, but also has to adapt to the different nutrient and ion availabilities, and avoid being out-competed or preyed on by the resident community^73^.

## Conclusions

This study represents the first comprehensive analysis of the evolution of saline and non-saline habitat preferences across the global tree of eukaryotes. We inferred that two of the largest assemblages of eukaryotes (TSAR and Amorphea) originated in different environments, and that ancestral eukaryotes likely inhabited non-marine environments. Our results show that marine and non-marine communities are phylogenetically distinct, but the salt barrier has been crossed several hundred times over the course of eukaryotic evolution. Several of these crossings coincided with the birth of diverse lineages, indicating that the availability of new niches has likely played a large role in the vast eukaryotic diversity we see today. We predict that the generation of genomic data from closely related marine and non-marine lineages will shed light on the genetic and cellular adaptations that have allowed crossings over the salt barrier.

## Methods

### Environmental samples for long-read metabarcoding and total DNA extraction

A total of 18 samples were sequenced for this study: five freshwater samples, four soil samples, four marine euphotic samples, and five marine aphotic samples (see Supplementary Table 1 for sample coordinates and details). Additionally, we used reads from three soil samples that were sequenced in a previous study^28^ (ENA accession PRJEB25197), resulting in a total of 21 samples that were analysed in this study. The aim here was to get a representative view of the microbial eukaryotic diversity in each environment.

#### Soil samples (x4 samples)

Peat samples were collected from (1) Skogaryd mire and (2) Kallkäls mire in October-November 2019. 5ml samples with three to four replicates of the top layer of soil were collected at both sites and visible roots were removed. Samples were kept at 4° for two days before extracting DNA using the DNeasy PowerSoil Kit (Qiagen). We also obtained DNA extracted from: (3) rainforest soil samples (six sites) from Puerto Rico ^75^, and (4) boreal forest soil samples (six sites) from Sweden^76^.

#### Freshwater samples (x5 samples)

We sampled three freshwater lakes in Sweden in October-November 2019: (1) Lake Erken, (2) Lake Ersjön, and (3) Lake Stortjärn. Plantonic samples were collected from the middle of the lakes at multiple depths, and mixed. Up to 3L of water was pre-filtered through a 200 μm mesh net to remove larger organisms before sequentially filtering through 20-25 μm, 3 μm, and 0.25 μm polycarbonate filters (47 mm). Filters were immediately frozen at -20° and stored at -70° before further processing. We also collected a (4) freshwater sediment sample (four replicates) from Lake Erken. The upper 0-5 cm of a sediment core was separated and mixed. All samples were kept at 4° before processing and extracting DNA using the DNeasy PowerSoil Kit. Lastly, we obtained DNA from (5) 10 permafrost thaw ponds in Canada^77^.

#### Marine euphotic samples (x4 samples)

One 5L sample was collected from the (1) North Sea at a depth of 5 m. Water was processed, and DNA extracted as described for the freshwater water samples. We used DNA extracts from the nano (3-20 μm) and pico (0.2-3 μm) fractions of two stations from the Malaspina expedition (Stations 49 and 76)^78^. These extracts corresponded to one (2) surface sample at 3 m depth, and (3-4) two DCM layer samples at depths of 70 m and 85 m.

#### Marine aphotic samples (x5 samples)

We used DNA extracts from the nano and pico fractions of the aphotic marine environment from Malaspina stations 49 and 76 ^78^. These corresponded to depths of (1-2) 275 m and 800 m for the mesopelagic, and (3-4) 1200 m and 2800-3300 m for the bathypelagic samples. Lastly, we obtained (5) DNA from a Mariana Trench sample from a depth of 5900 m^79^.

### PCR amplification and long-read sequencing

We amplified a ∼4500 bp fragment of the ribosomal DNA operon, spanning the 18S gene, ITS region, and 28S gene, using the general eukaryotic primers 3NDf ^80^ and 21R^81^. PCRs were performed with sample-specific tagged-primers using the Takara LA Taq polymerase (Takara) and 5 ng of DNA as input. PCR-cycling conditions included an initial denaturation step at 94° for 5 min, at least 25 cycles of denaturation at 98° for 10 sec, primer annealing at 60° for 30 sec, and elongation at 68° for 5 min, and finishing with a final elongation step at 68° for 10 min. We limited the number of PCR cycles to 25, where possible, to reduce chimera formation^82^. For samples that did not get amplified, we increased the number of cycles to 30. PCR products were assessed using agarose gels and Qubit 2.0 (Life Technologies), and then purified with Ampure XP beads (Beckman Coulter). Amplicons from replicates and different sites from the same sampling location were pooled at this stage. SMRTbell libraries were constructed using the HiFi SMRTbell Express Template Prep Kit 2.0. Long-read sequencing was carried out at SciLifeLab (Uppsala, Sweden) on the Sequel II instrument (Pacific Biosciences) on a SMRT Cell 8M Tray (v3), generating four 30-hour movies.

### Processing reads and OTU clustering

We QC filtered sequences following^28^ with some modifications. The CCS filtration pipeline is available at ^83^. Briefly, Circular Consensus Sequences (CCS) were generated by SMRT Link v8.0.0.79519 with default options. The CCS reads were demultiplexed with mothur v1.39.5^84^, and then filtered with DADA2 v1.14.1^85^. Reads were retained if they had both primers and if the maximum number of expected errors was four (roughly translating to one error for every thousand base pairs). We pre-clustered reads at 99% similarity using VSEARCH v2.3.4^86^, and generated consensus sequences for pre-clusters ≥ 3 reads to denoise the data. Prokaryotic sequences were detected by BLASTing^87^ against the SILVA SSU Ref NR 99 database v132^88^ and removed. We predicted 18S and 28S sequences in the reads using Barrnap v0.7 (--reject 0.4 --kingdom euk)^89^, and discarded non-specific and artefactual reads (i.e. those containing multiple 18S/28S, or missing 18S/28S). Chimeras were detected *de novo* using Uchime^90^ as implemented in mothur. Finally, we extracted the 18S and 28S sequences from the reads and clustered them using VSEARCH into Operational Taxonomic Units (OTUs) at 97% similarity. After discarding singletons, a second round of *denovo* chimera detection was performed using VSEARCH, and chimeric OTUs were removed. We calculated sequence similarity of the OTUs against reference sequences in a custom PR^2^ database^30^ (*PR2-transitions*^31^; see below) using VSEARCH (--usearch_global and --iddef 1). All references and OTU sequences were trimmed with the primers 3ndf and 1510R^91^ to ensure that they spanned the same region.

### Taxonomic annotation of long-read sequences

#### The modified PR2 reference database

Reference sequences were derived from a modified version of the Protist Ribosomal Reference (PR2) database v4.12.0^30^, called *PR2_transitions*. This database revised the taxonomy structure of PR2 to 9 levels: Domain, Supergroup, Division, Subdivision, Class, Order, Family, Genus, Species. This allowed us to update the taxonomy to accommodate recent changes in eukaryotic classification^92^ (changes in taxonomy can be viewed at ^83^). Additionally, we added sequences from nucleomorphs, and several newly discovered or sequenced lineages such as Rholphea, Hemismastigophora, and others. *PR2_transitions* is available on Figshare^31^. We used the 18S gene alone for taxonomic annotation, as 28S databases are much less comprehensive by comparison.

#### Phylogeny-aware taxonomy assignment

We used a phylogeny-aware approach to assign taxonomy to the PacBio OTUs, as done in^28^. This approach assigns taxonomy to the appropriate taxonomic rank, such that OTUs branching deep in the eukaryotic tree are labelled to high taxonomic ranks, and vice versa. For each sample, we inferred preliminary maximum likelihood trees along with SH-like support^93^ with RAxML v8^32^ (using the GTRCAT approximation as it is better suited for large trees^94^). These trees contained the filtered OTUs and closely related reference sequences from *PR2_transitions*. Trees were scanned manually to identify mis-annotated reference sequences, nucleomorphs, and artefactual OTUs. After removing these sequences, we inferred trees with RAxML-NG^95^ using 20 starting trees.

The final taxonomy was generated by getting the consensus of two strategies. Strategy 1 parses the tree and propagates taxonomy to the OTUs from the nearest reference sequences using the Genesis^96^ app *partial-tree-taxassign*^97^. Strategy 2 starts by pruning the OTUs from the phylogeny, leaving behind references only. OTUs are then phylogenetically placed on the tree with EPA-ng v0.3.5 ^34^, and taxonomy assigned using the gappa^96^ command *assign* under the module *examine*. The resulting taxonomy of the 18S gene of each OTU was transferred to its 28S gene counterpart, as the molecules are physically linked.

### Maximum likelihood analyses of the global eukaryotic dataset

18S and 28S sequences were aligned using MAFFT v7.310^98^ using the FFT-NS-2 strategy, and subsequently trimmed with trimAl^99^ to remove sites with >95% gaps. We inferred preliminary trees from a concatenated alignment with RAxML v8.2.12 under the GTRCAT model^32^ which were then visually inspected to detect chimeras and sequence artefacts. Taxa were removed if their position in the tree did not match their taxonomy. Four such rounds of visual inspection were performed, two with unconstrained trees, and two with constrained trees (see text below for details on constraints). To avoid long branch attraction, we excluded rapidly evolving taxa using TreeShrink^100^ (k=2500). This resulted in the removal of *Mesodinium*, long-branch Microsporidia, several Apicomplexa, several Heterolobosea, and several Colladaria from our dataset.

After removing chimeras and sequence artefacts, we realigned and trimmed the 18S and 28S sequences as before. After concatenation, the final dataset was composed of 16,821 taxa and 7,160 alignment sites. Global eukaryotic phylogenies of the taxonomically annotated, 18S-28S environmental sequences were inferred using RAxML v8.2.12 under the GTRCAT model^32^, and 100 transfer bootstrap replicates (TBE)^101^. Supergroups, Divisions, and Subdivisions (ranks 2, 3 and 4 in *PR2_transitions*) were constrained to be monophyletic in our tree (i.e. all taxa labelled as a specific subdivision were constrained to be on one side of a split). The one exception was Excavata whose monophyly has not been confidently resolved^29^. One hundred maximum likelihood inferences were performed in order to take phylogenetic uncertainty into account for subsequent ancestral state reconstruction analyses. We opted to include only the long-read environmental sequences in our phylogenies because they better represent environmental diversity (compared to reference databases which are more biased towards culturable organisms and marine environments^102^), and because very few 18S-28S sequences can otherwise be ascertained to derive from the same organism. The final tree along with metadata was visualised using the anvi’o interface^103^ and then modified in Adobe Illustrator^104^ to label clades.

### Short read datasets

#### Datasets collected

Short-read data corresponding to the V4 hypervariable region were retrieved from 22 publicly available metabarcoding datasets. Data were considered if the following criteria were fulfilled: (i) samples were collected from soils, freshwater, or marine habitats (ii) there was clear association between samples and environment (i.e. no data from estuaries where salinity fluctuates); and (iii) data publicly available or authors willing to share. The search for studies was not meant to be exhaustive and the datasets included in this work were identified and collected by the end of October 2020, unless specified otherwise. A list of these datasets can be found in Supplementary Table 3.

#### Processing short-read data and clustering into OTUs

Raw sequence files and metadata were downloaded from NCBI SRA web site^105^ when available or obtained directly from the investigators. Information about the study and the samples (substrate, size fraction etc.) as well as the available metadata (geographic location, depth, date, temperature etc.) were stored in three distinct tables in a custom MySQL database stored on Google Cloud. For each study, raw sequences files were processed independently de novo. Primer sequences were removed using cutadapt^106^ (maximum error rate = 10%). Amplicon processing was performed under the R software^107^ using the dada2 package^85^. Read quality was visualized with the function *plotQualityProfile*. Reads were filtered using the function *filterAndTrim*, adapting parameters (truncLen, minLen, truncQ, maxEE) as a function of the overall sequence quality. Merging of the forward and reverse reads was done with the *mergePairs* function using the default parameters (minOverlap = 12, maxMismatch = 0). Chimeras were removed using *removeBimeraDenovo* with default parameters. Taxonomic assignation of ASVs was performed using the *assignTaxonomy* function from dada2 against the PR2 database^30^ version 4.12 (https://pr2-database.org). ASV assignation and ASV abundance in each sample were stored in two tables in the MySQL database. ASV information was retrieved from the database using an R script.

ASVs from each environment (freshwater, soil, marine euphotic, marine aphotic) were clustered into OTUs at 97% similarity using VSEARCH^86^, to make the size of the dataset more manageable for subsequent phylogenetic analyses. To be conservative in what was considered to be present in an environment, we retained only those OTUs that were composed of at least 100 reads, or were present in multiple distinct samples.

### Phylogenetic placement on global eukaryote phylogeny

Short-read OTUs were aligned against the long-read alignment (see **Maximum likelihood analyses of the global eukaryotic dataset**) using the phylogeny-aware alignment software PaPaRa^108^. Misaligned sequences were systematically checked and removed. OTUs from the four environments were then phylogenetically placed on the global eukaryote tree (the tree with the highest likelihood) using EPA-ng^34^. OTUs with high EDPL (expected distance between placement locations) indicate uncertainty in placement, and were filtered out with the gappa command *edpl*^96^. The resulting jplace files were visualised with iTOL^109^.

### Inferring clade specific phylogenies with short- and long-read data

In order to investigate clade-specific transition rates across the salt barrier, we inferred phylogenies for major eukaryotic groups. We considered only those clades that contained sufficient data to more precisely infer transition rates; i.e. both terrestrial and marine taxa were present, and there were at least 50 taxa present. This excluded taxa such as radiolarians (which contains no terrestrial taxa), rigifilids (which contains only terrestrial taxa), and tubulineans (which is predominantly terrestrial with an extremely small proportion of marine taxa). After preliminary analyses, we also excluded the clades discobans and discoseans due to large topological differences in the resulting trees.

We extracted all short-read OTUs from the remaining 13 clades using the gappa subcommand *extract*. Short-read OTUs taxonomically annotated as anything other than the respective clade were discarded (for instance we discarded sequences labelled as amoebozoans that were phylogenetically placed in apicomplexa). For each clade, we pruned the corresponding subtree (and an outgroup) from the global phylogeny with the best likelihood score. For each clade, we then inferred 100 ML phylogenies with RAxML (GTRCAT model), using the long-read subtree as a backbone constraint.

### Analyses of habitat evolution

#### Unifrac analyses

To estimate whether microbial communities from various habitats were phylogenetically distinct, we calculated unweighted UniFrac distance^110^ as implemented in mothur, between (1) marine and terrestrial habitats, (2) marine euphotic, marine aphotic, soil, and freshwater, and (3) each sample sequenced with PacBio. Distances were estimated along the best ML global eukaryotic phylogeny with 1000 randomisations in order to test for statistical significance.

Similarly, we estimated pairwise Kantorovich-Rubinstein distance (earth mover’s distance) between the four habitats (soil, freshwater, marine euphotic, marine aphotic) using the gappa subcommand *krd* with the short-read placement files (jplace files) as input (See **Phylogenetic placement on global eukaryotic phylogeny**).

#### Model test on global eukaryotic phylogeny

To investigate whether transition rates vary between major eukaryotic clades, we compared a null model (qMT and qTM remain constant throughout the global eukaryotic tree) against a complex model (qMT and qTM estimated separately for each major eukaryotic clade) on the global eukaryotic phylogeny. These models were compared using MCMC analyses in BayesTraits v3.0.2^111,112^ in a reversible-jump framework in order to avoid over-parameterization^113^. Following the analysis in ^114^, we used 50 stones and a chain length of 5,000 to obtain marginal likelihood for each model using stepping stone method^115^, and a Log Bayes Factor (2 * difference of log marginal likelihoods) of 10 or more was used to favour the complex model over the simple model.

Before final analyses in BayesTraits, we tried several prior distributions for transition rates (using a hyperprior approach to reduce uncertainty about prior choice^113^). Specifically we compared gamma hyperpriors with exponential hyperpriors using different values. While the different priors produced qualitatively similar results, we found the exponential hyperprior to be most suitable. All BayesTraits analyses were therefore carried out using an exponential hyperprior with the mean seeded from a uniform distribution between 0 and 2. Additionally, all ancestral state reconstruction analyses were carried out on 100 inferred phylogenies to take phylogenetic uncertainty into account, and were repeated thrice to check for convergence.

#### Clade specific transition rates

We inferred clade-specific transition rates along the clade-specific phylogenies (long-read + short-read data), on account of these being more complete. The metadata for each taxon was used to label it as either marine or terrestrial. We ran 1 million generations on each tree (100 million generations in total) with 0.5 million generations discarded as burn-in. For each clade, we also inferred the global transition rate, regardless of the direction of transition. This was achieved by normalising the QMatrix^116,117^, with all other parameters unchanged. These analyses also allowed us to infer the ancestral state of each major eukaryotic clade.

#### Inferring ancestral states of deep nodes and the last common ancestor of eukaryotes

In order to infer the ancestral habitats at deeper nodes (including the origin of eukaryotes), we modelled habitat evolution along the global eukaryotic phylogeny using the better suited complex model. Analyses were run for 500 million generations, forcing BayesTraits to spend 5 million generations on each tree, and 200 million generations were discarded as burn-in. Analyses were carried out after rooting the tree at Discoba, and at Amorphea in order to take uncertainty about the root into account.

#### Visualising scenarios of habitat evolution

Most ancestral state reconstruction programmes do not explicitly calculate the ancestral state at internal nodes (but integrate over all possibilities). In order to visualise habitat evolution, we used PastML, a maximum likelihood ancestral state reconstruction programme which calculates the state at each internal node, and also generates a concise visual summary of the clade. For each major eukaryotic clade, we ran PastML on 100 trees. Visualisations for several trees were checked manually to assess if they displayed similar histories, and one visualisation was chosen randomly for display in Supplementary Figure 12.

#### Counting number and relative timing of transitions

We converted all clade-specific phylogenies into relative chronograms (with the age of the root set to 1) using Pathd8^38^ which is suitable for large phylogenies. We ran PastML on these phylogenies (as before), and used custom scripts^83^ to count the number of marine-terrestrial transitions. For each transition, we calculated the distance to the root to obtain relative timing of transition.

### Network analyses

To check that our results about transition rates and timings were not biased by phylogenetic inference from sequences with poor phylogenetic signal, we constructed sequence similarity networks. These networks were constructed using representative 18S sequences of the long-read OTUs. Briefly, we performed all-against-all BLAST searches, and generated networks using a coverage threshold of 75, and sequence identity thresholds of 80, 85, 90, 95, 97. Networks were visualized on Cytoscape^118^. Assortativities were calculated using scripts available at ^119^, and then plotted in R using *ggplot*^120^.

### Data availability

New sequence data generated for this study were deposited at ENA under the accession number PRJEB45931, while data from Sequel I (generated in ^28^) were deposited under the accession number PRJEB25197. The *PR2-transitions* database, annotated 18S and 28S OTU sequences, clustered short read metabarcoding sequences used in this study, and all trees have been deposited in an online repository^31^. All custom code is available here^83^.

## Acknowledgments

We want to thank Anna Rosling, Hector Urbina, and Matias Cafaro for kindly providing DNA from soil samples collected in Sweden and Puerto Rico. We thank the pilots of the deep-sea HOV “Jiao Long Hao” and the crew of the R/Vs “Xiang Yang Hong 09” for their professional service during cruise DY37II to collect samples from the Mariana Trench. We are grateful to the Swedish Infrastructure for Ecosystem Science (SITES) for collecting samples from Swedish lakes, and Swedish Meteorological and Hydrological Institute (SMHI) for collecting a sample from the North Sea. Marine sampling was supported by the Spanish Ministry of Economy, Competitiveness projects Malaspina-2010 (CSD2008–00077) and ALLFLAGS (CTM2016-75083-R). We would like to thank Éléna Coulier for her help with optimising the long-range PCRs. We thank Olga V. Petterson and Christian Tellgren-Roth for designing fusion primers for long-read amplification. We thank Miguel M. Sandin for his help with network analyses, and Jesper Boman for help with awk scripting. We thank Javier del Campo for his advice on updating taxonomy for our custom PR2 database. We thank the ABIMS platform of FR2424 (CNRS, Sorbonne Université) for bioinformatics resources. The authors would like to acknowledge support of the National Genomics Infrastructure (NGI) / Uppsala Genome Center and UPPMAX for providing assistance in massive parallel sequencing and computational infrastructure (SNIC 2021/5-302). Work performed at NGI / Uppsala Genome Center has been funded by RFI / VR and Science for Life Laboratory, Sweden. This work was supported by a grant from Science for Life Laboratory available to F.B., which covered the salary of M.J., and experimental expenses.

## Author contributions

F.B. and M.J. conceived the project. M.J., H.J., S.P., and R.M. collected samples and extracted DNA. M.J. carried out long-range PCRs and processed the PacBio data. C.B. and D.V. collected and processed short-read metabarcoding data. A.O. performed comparisons of long and short-read metabarcoding data. M.J, and C.B. performed phylogenetic and ancestral state reconstruction analyses. M.J and F.B. wrote the first draft of the manuscript and all authors read and commented on the manuscript. F.B. supervised the project.

