## Supplementary Figures and Tables for "Global patterns and rates of habitat transitions across the eukaryotic tree of life"

### Supplementary Information

Mahwash Jamy[1], Charlie Biwer[1], Daniel Vaulot[2,3], Aleix Obiol[4], Homgmei Jing[5], Sari Peura[6,7], Ramon Massana[4], Fabien Burki[1,7*]

[1] Department of Organismal Biology (Systematic Biology), Uppsala University, Uppsala, Sweden

[2] Sorbonne Université, CNRS, UMR7144, Team ECOMAP, Station Biologique, Roscoff, France

[3] Asian School of the Environment, Nanyang Technological University, Singapore

[4] Department of Marine Biology and Oceanography, Institut de Ciències del Mar (ICM-CSIC), Barcelona, Spain

[5] CAS Key Lab for Experimental Study Under Deep-sea Extreme Conditions, Institute of Deep-sea Science and Engineering, Chinese Academy of Sciences, Sanya, China

[6] Department of Ecology and Genetics (Limnology), Uppsala University, Uppsala, Sweden

[7] Science for Life Laboratory, Uppsala University, Sweden

**a**

| | reads | | | ASVs | | | Total | |
|---|---|---|---|---|---|---|---|---|
| Dataset | 49/DCM | 49/Meso | 76/Surf | 49/DCM | 49/Meso | 76/Surf | reads | ASVs |
| mTags | 846 | 822 | 470 | - | - | - | 2138 | - |
| V4 | 77451 | 58953 | 147493 | 1740 | 1183 | 1800 | 283897 | 3753 |
| V9 | 46004 | 23642 | - | 2034 | 1127 | - | 69646 | 2777 |
| PacBio | 273073 | 642804 | 149790 | 4728 | 2277 | 1357 | 1065667 | 8362 |

**b**

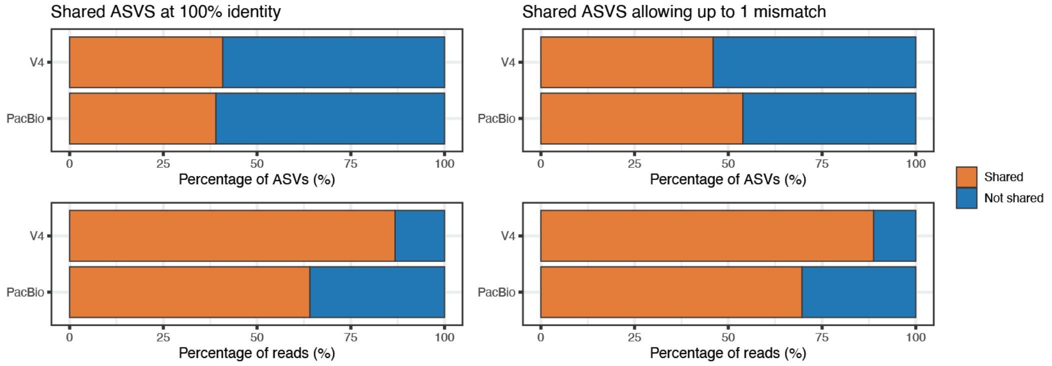

**Supplementary Figure 1.** Comparison of PacBio and Illumina sequencing. PacBio amplicons were compared with metagenomes (mTags), V4 amplicons, and V9 amplicons from three marine samples corresponding to the pico size fraction from the Malaspina expedition[1]. Station 76|Surface did not have V9 amplicon data. ASVs = Amplicon Sequence Variants.

**(a)** Number of reads and ASVs for each sample for each marker. The mTags represent sequence length of ca. 100 bp, so no ASV level is available, as this short length does not give enough resolution. More PacBio sequences were generated for each sample compared to Illumina sequences.

**(b)** Comparison of PacBio ASVs (i.e. de-noised, preclustered sequences) with the ones given by V4 amplicons. A similar comparison with V9 ASVs was not carried out as not all samples had V9 Illumina data available. Around half of the sequences were shared, which represented the majority of reads.

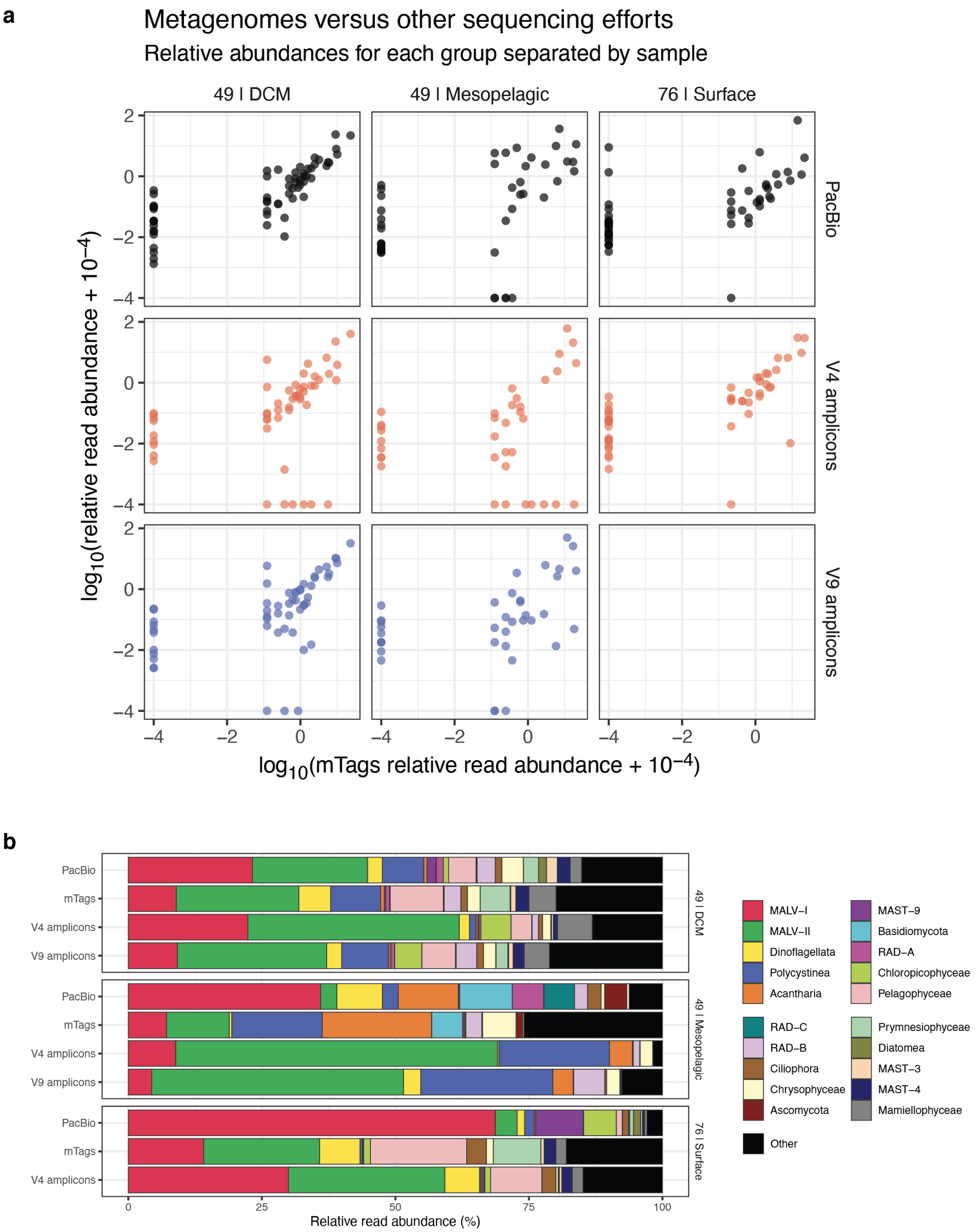

**Supplementary Figure 2.** Comparison of the eukaryotic communities retrieved by PacBio and Illumina sequencing (V4, V9, and 18S reads retrieved from metagenomic data) of three marine samples (See Supplementary Figure 1).

**(a)** Comparison of mTags (which should represent a snapshot of the community unbiased by PCR) with the other datasets. Groups explaining the majority of reads are detected at comparable abundances. Points at the margins represent taxa that are found in one dataset but not in the other;

along the x axis we see groups that are present in mTags but not in the other datasets. For instance in the 49|DCM panel, there are some groups in mTags that V4/V9 amplicons cannot detect (blue and red points at the bottom). The line of dots along the y-axis represent groups not present in mTags, but present in other datasets. Fewer black points (PacBio) at the bottom of the panels, indicates that PacBio is detecting groups that are missed by metabarcoding with V4/V9 sequencing.
**(b)** Overall comparison of the relative abundances at the group level (excluding Charophyta, Metazoa and Nucleomorphs). The primer pair used for long-read sequencing seem to preferentially amplify MALV-I, but the overall community structure that PacBio is retrieving is reasonable with the other sequencing approaches.

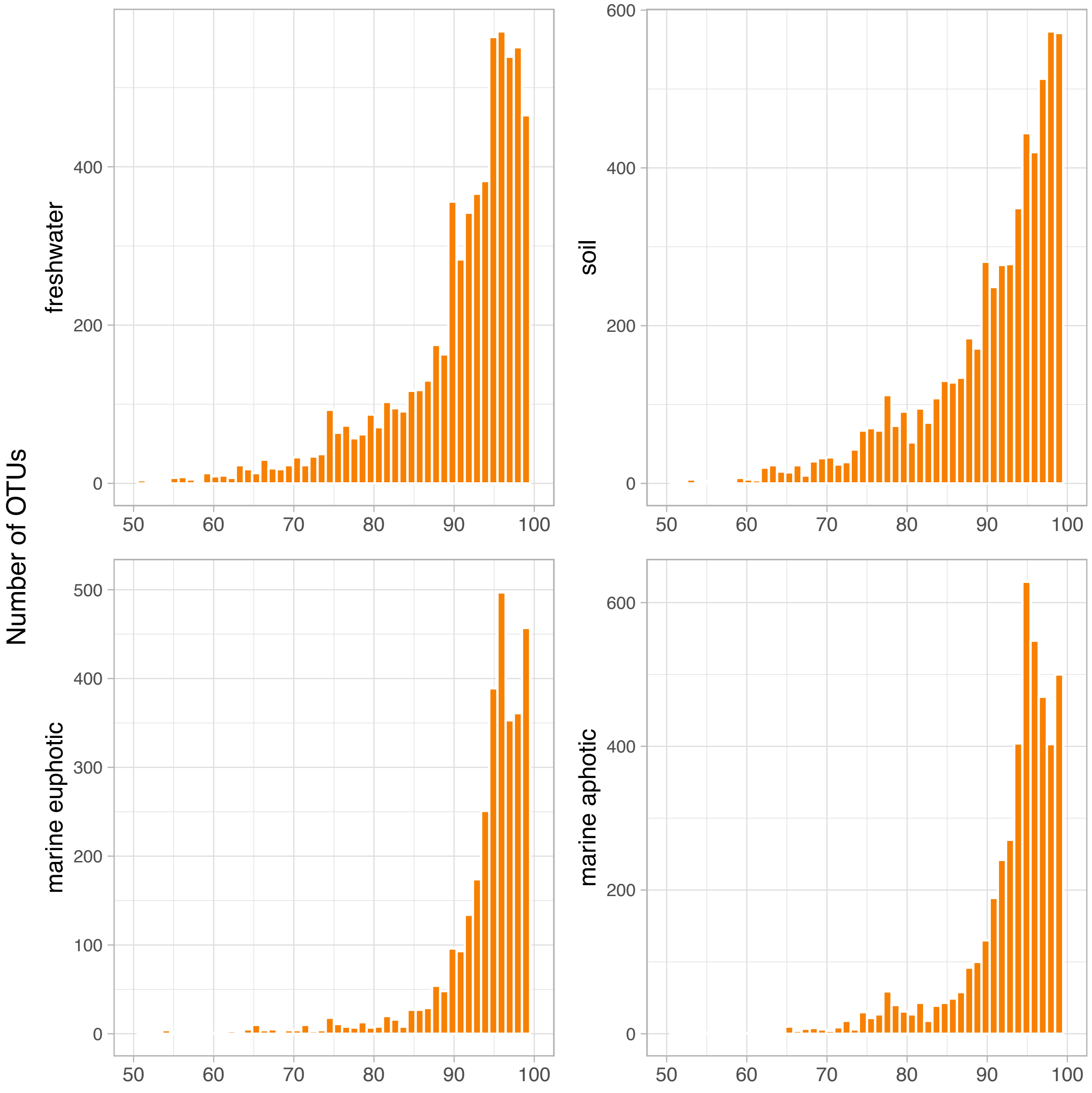

**Supplementary Figure 3.** Percentage similarity of OTUs (18S sequence only) against reference sequences in the PR2 database[2], as determined by vsearch global search[3]. All sequences (queries and references) were trimmed with primers 3NDF and 1510R so that they spanned the same region.

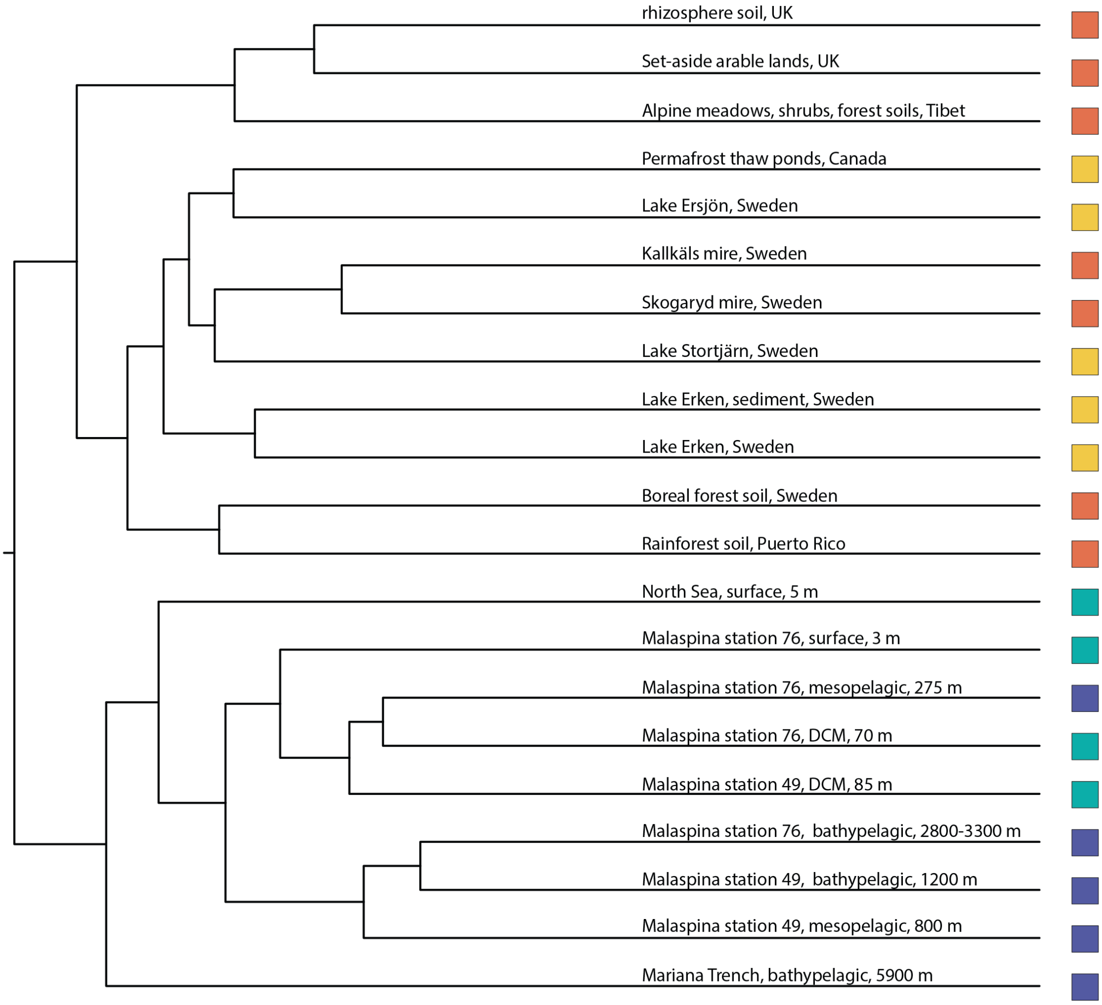

**Supplementary Figure 4.** Hierarchical clustering of the 21 samples sequenced with PacBio based on unweighted UniFrac analyses on the tree in Figure 1a. UniFrac measures the dissimilarity between communities by taking phylogenetic relatedness into account[4]. Here, soil=orange, yellow, freshwater, light blue=marine photic, and dark blue=marine aphotic samples. The figure shows a deep split between marine and terrestrial communities. Furthermore, on average, marine communities seem to be more closely related to each other than terrestrial communities.

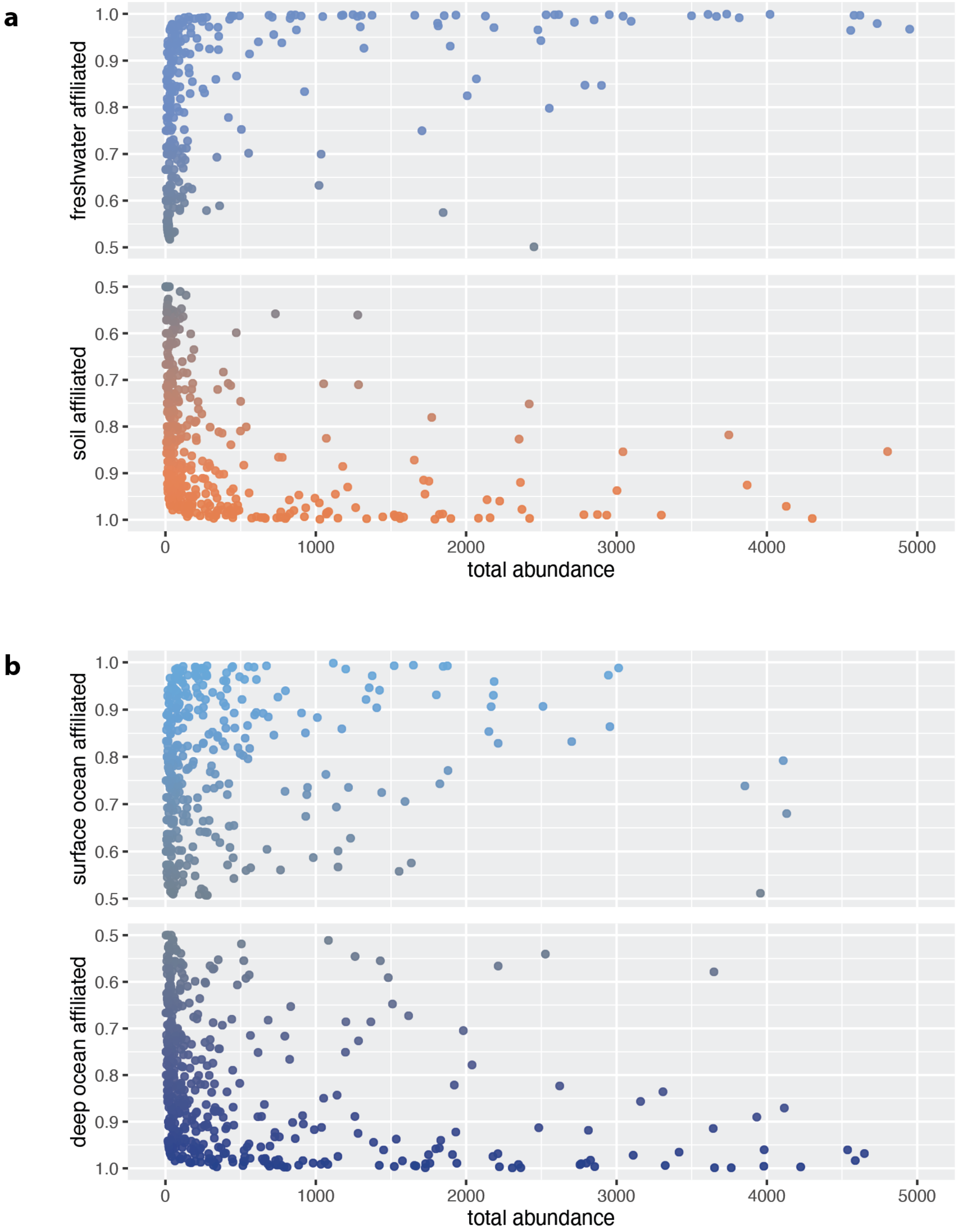

**Supplementary Figure 5.** Distribution patterns of **(a)** the 723 (out of 771) shared OTUs between freshwater and soil PacBio samples, and **(b)** the 801 (out of 854) shared OTUs between the surface and deep ocean. All OTUs were clustered at 97% similarity. The x-axis represents the total number of reads in the OTU (a cut-off of 5000 reads was chosen as shared OTUs with higher abundances were spurious), and the y-axis represents affiliation to each environment as measured by the (number of reads in a habitat) / (total number of reads). This analysis follows the reasoning presented in [5]. Briefly, shared OTUs do not necessarily represent generalists across two habitats, but can also represent contamination, i.e. taxa that ended up in another habitat due to run-off, water mixing etc. To differentiate between the two, the affiliation to each habitat can be measured. For instance, if a shared OTU between soil and freshwater habitats has roughly half of its reads present in soil and half in freshwater, then it can be assumed that the taxon is more likely to be a generalist. However, if the shared OTU has 90% of its reads in soil habitats, and only 10% in freshwater, it is more likely that the taxon lives in soil, but ended up in freshwater due to run-off or some other way. The analysis shows a number of generalists in both **(a)** and particularly in **(b)**. As in Sieber et al. [5], “generalists” tend to be taxon with lower read abundances overall. Since the OTUs were clustered at 97% similarity, it is also possible that “generalists” are represented by closely related species with their own specific niches.

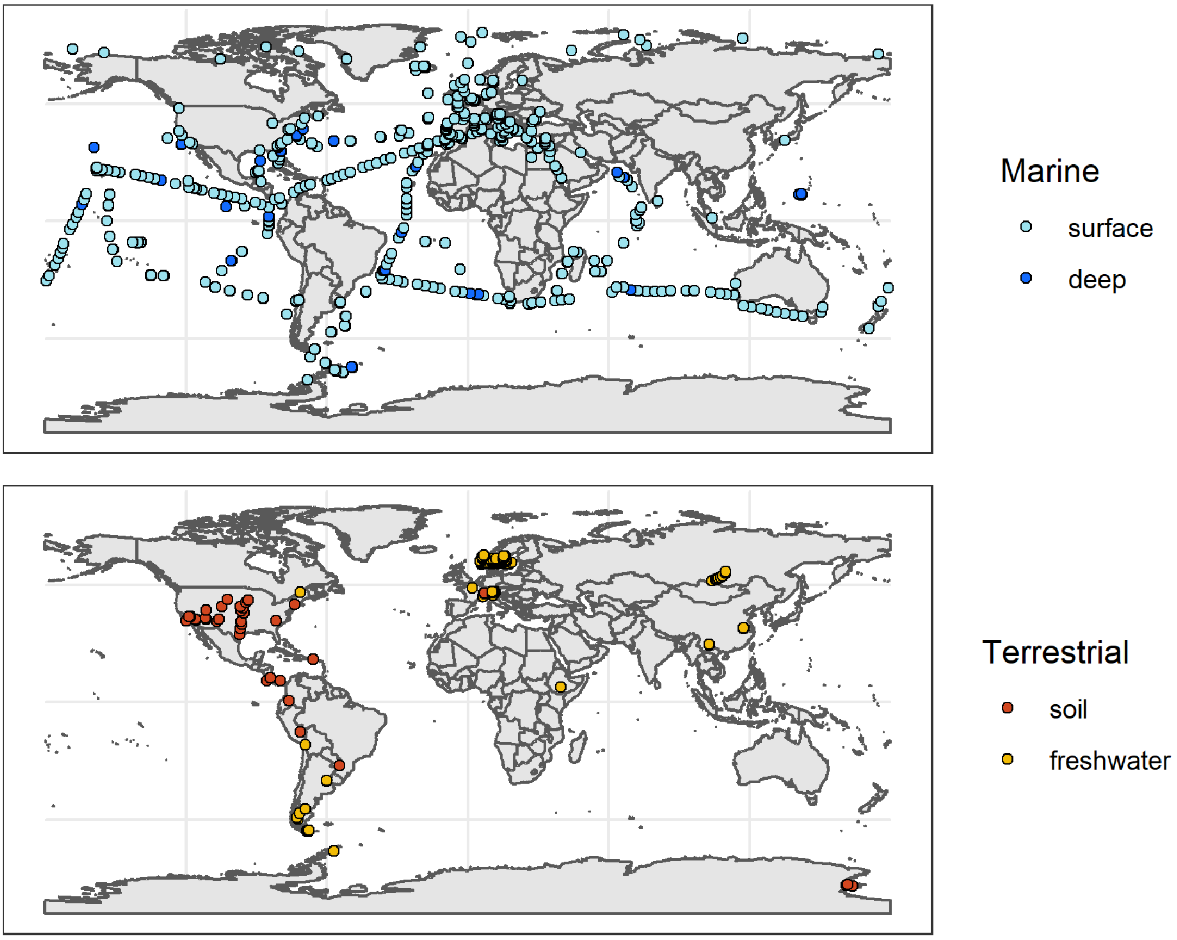

**Supplementary Figure 6.** Geographical locations corresponding to the short-read metabarcoding data. The upper panel showcases the provenance of marine samples, from the marine euphotic (light blue) and the marine aphotic (dark blue). The lower panel displays samples originating from terrestrial habitats, with soil samples in red and samples from freshwater systems (lakes, ponds, creeks and rivers) in yellow.

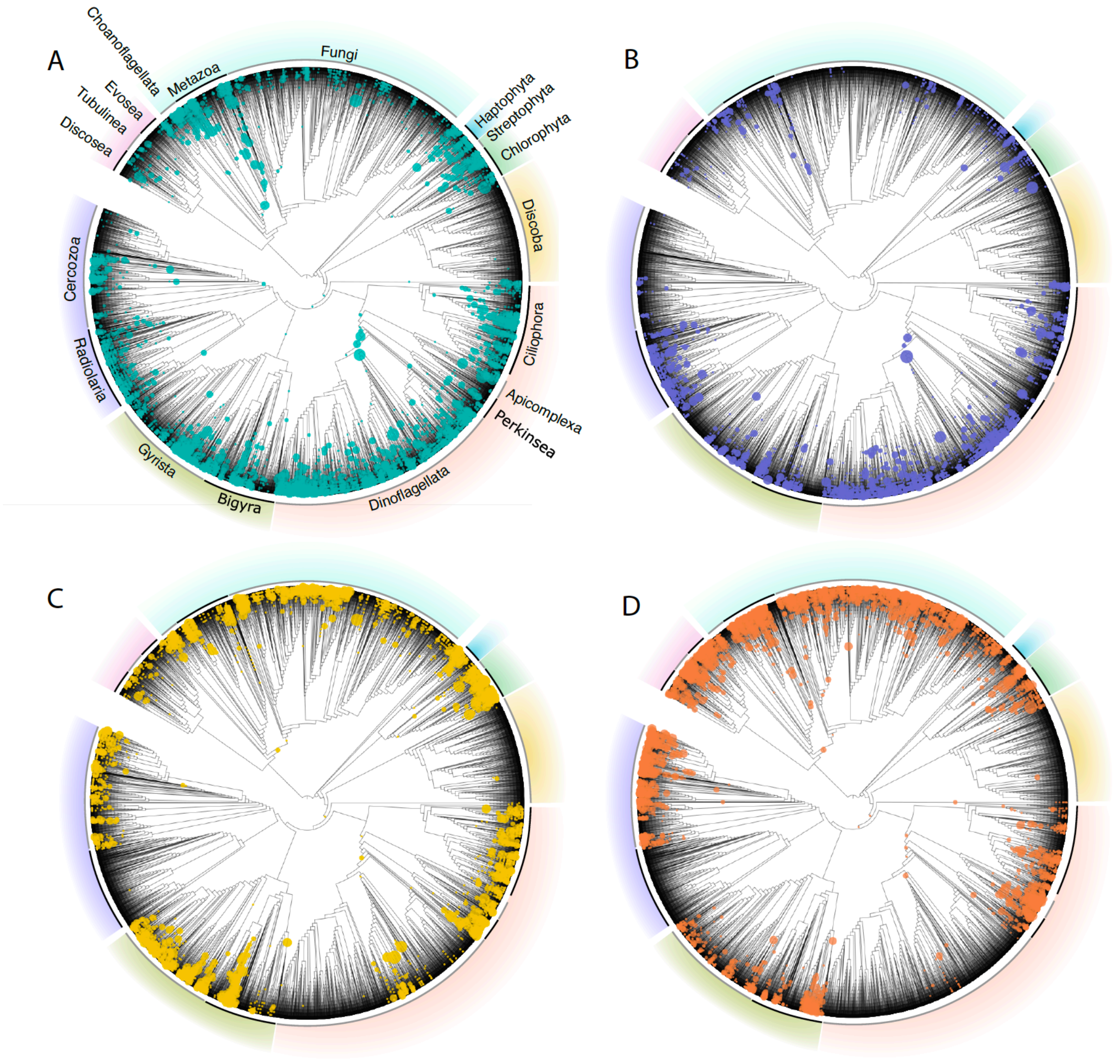

**Supplementary Figure 7.** Phylogenetic placement of short-read OTUs onto the long-read, global eukaryotic reference phylogenetic tree (in Figure 1). The upper two panels represent marine environments (A, marine euphotic; B, marine aphotic), while the lower two panels showcase terrestrial placements (C, freshwater; D, soil). Visualisation of the placement files was done through the interactive Tree of Life[6], and the size of each circle represents the number of placements on that particular branch weighted by the likelihood weight ratios.

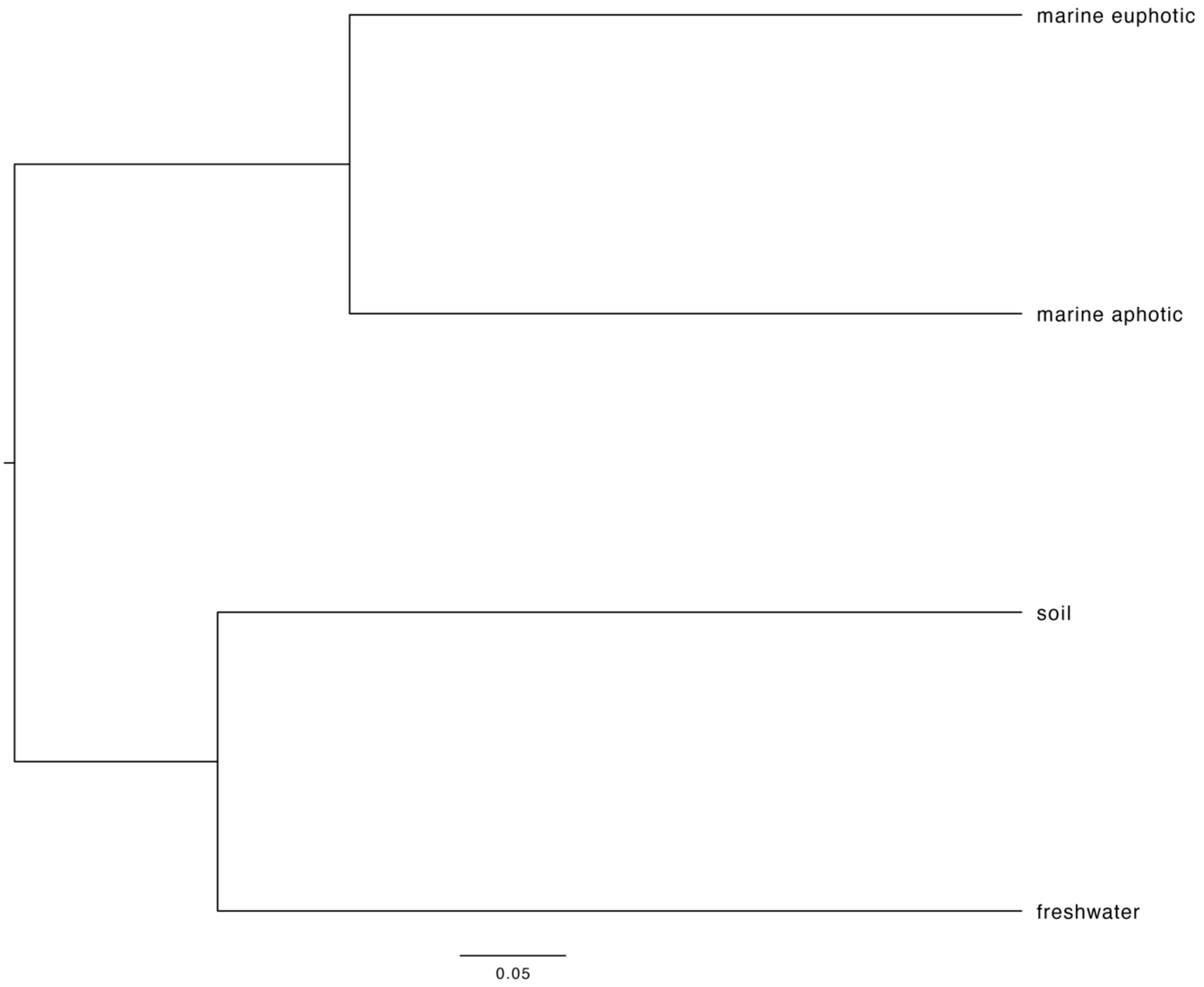

**Supplementary Figure 8.** UPGMA dendrogram based on Earth mover's distance between placement distributions of marine euphotic, marine aphotic, soil, and freshwater short-read OTUs on the global eukaryotic long-read phylogeny (Supplementary Figure 7). The earth mover's distance in this context refers to the minimum amount of "work" required to shift the placement distribution of one habitat to the distribution of another habitat.

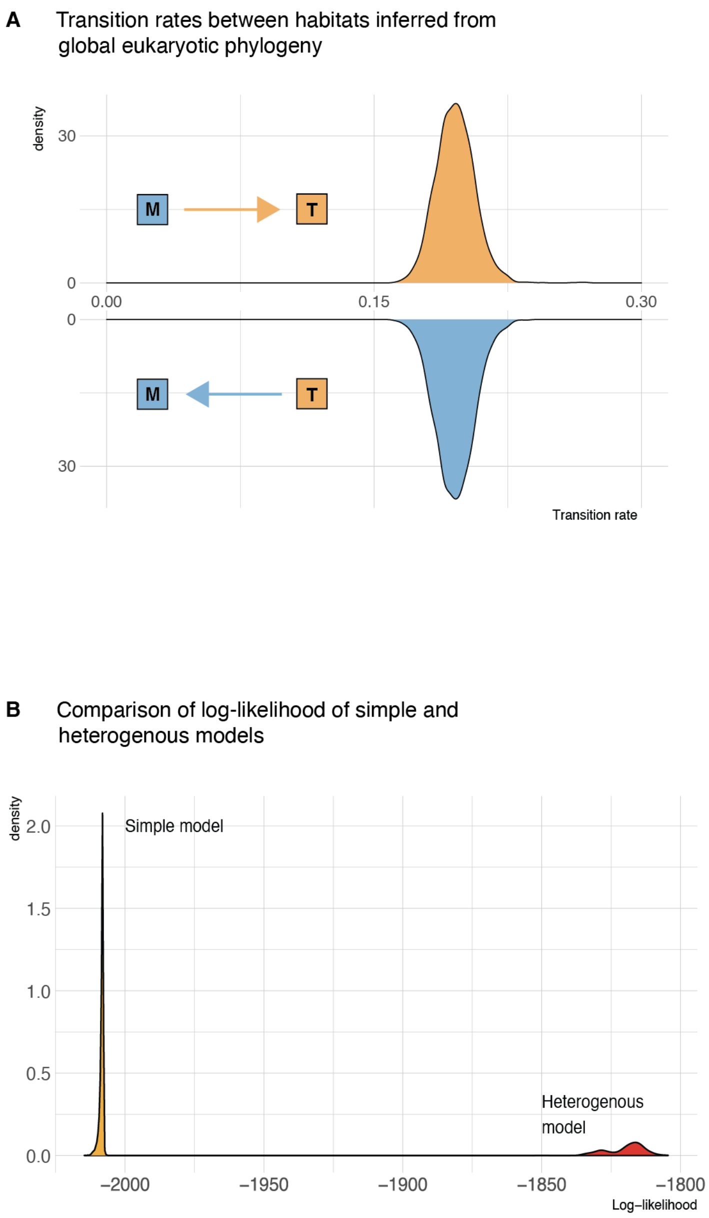

**Supplementary Figure 9. (A)** Instantaneous transition rates from marine to terrestrial habitats (qMT) and vice versa (qTM) when using a homogenous model over the global eukaryotic phylogeny (in Figure 1). This homogenous model allowed qMT and qTM to be unequal, but does not allow qTM and qMT to vary over the tree. **(B)** Comparison of the posterior probability of log-likelihoods when using the simple, homogenous model and the heterogenous model. The heterogenous model estimated a separated qMT and qTM for every major eukaryotic lineage (defined in this paper as rank 4 in the PR2-transitions database, e.g. Ciliates, Dinoflagellates, Fungi, etc.) that had at least 50 taxa and contained both marine and terrestrial taxa. The plot shows that the heterogenous model had a much better fit, indicating that rates of habitat evolution vary strongly across the eukaryotic tree of life.

A

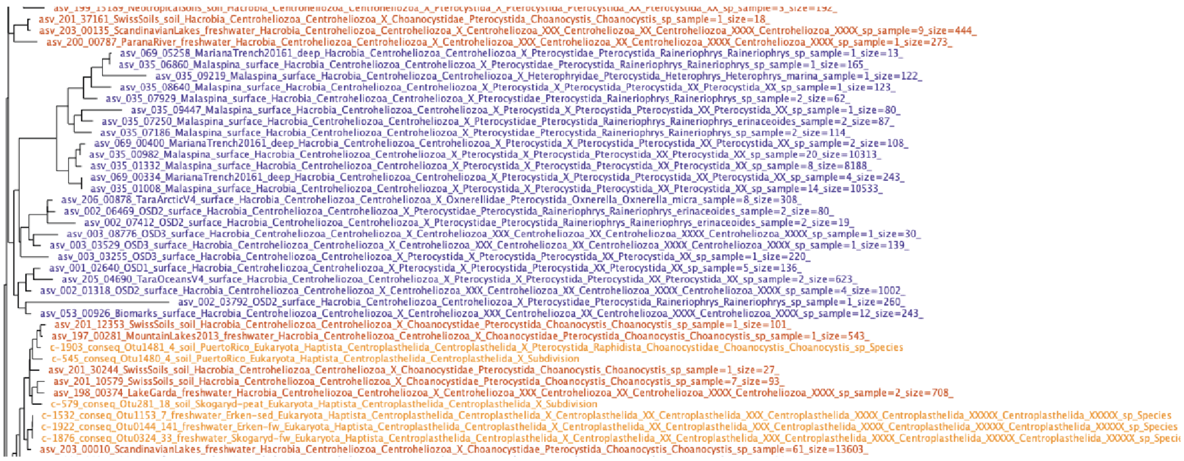

B

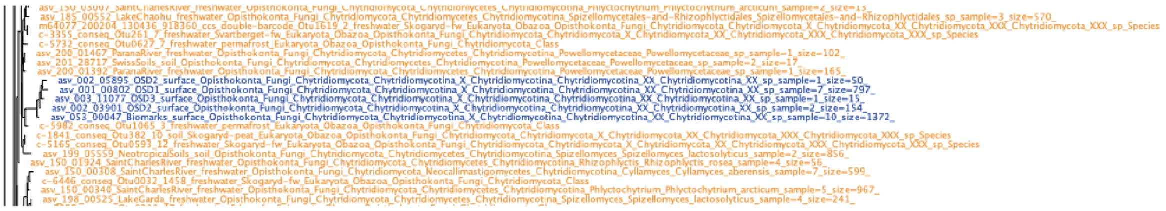

C

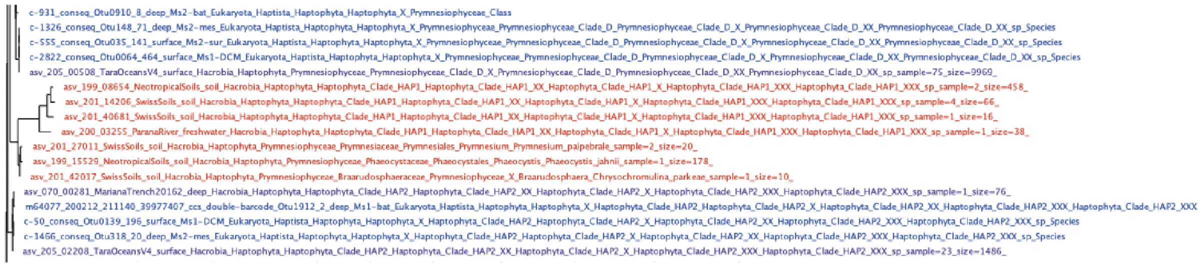

**Supplementary Figure 10.** Three examples of transitions detected by the incorporation of short-read data in our phylogenies that would otherwise have been missed. Shades of blue/purple represent marine sequences, while shades of orange/red represent terrestrial taxa. **(A)** A clade of marine Centroheliozoans is detected in purple (including sequences from the Malaspina expedition, Ocean Sampling Day, Tara Oceans, and Mariana Trench datasets). **(B)** A clade of marine Chytrids is detected mainly from Ocean Sampling Day datasets. **(C)** A clade of terrestrial Haptophytes is detected mainly from the Swiss Soils and Neotropical soil datasets. Such cases were spread throughout the eukaryotic phylogeny.

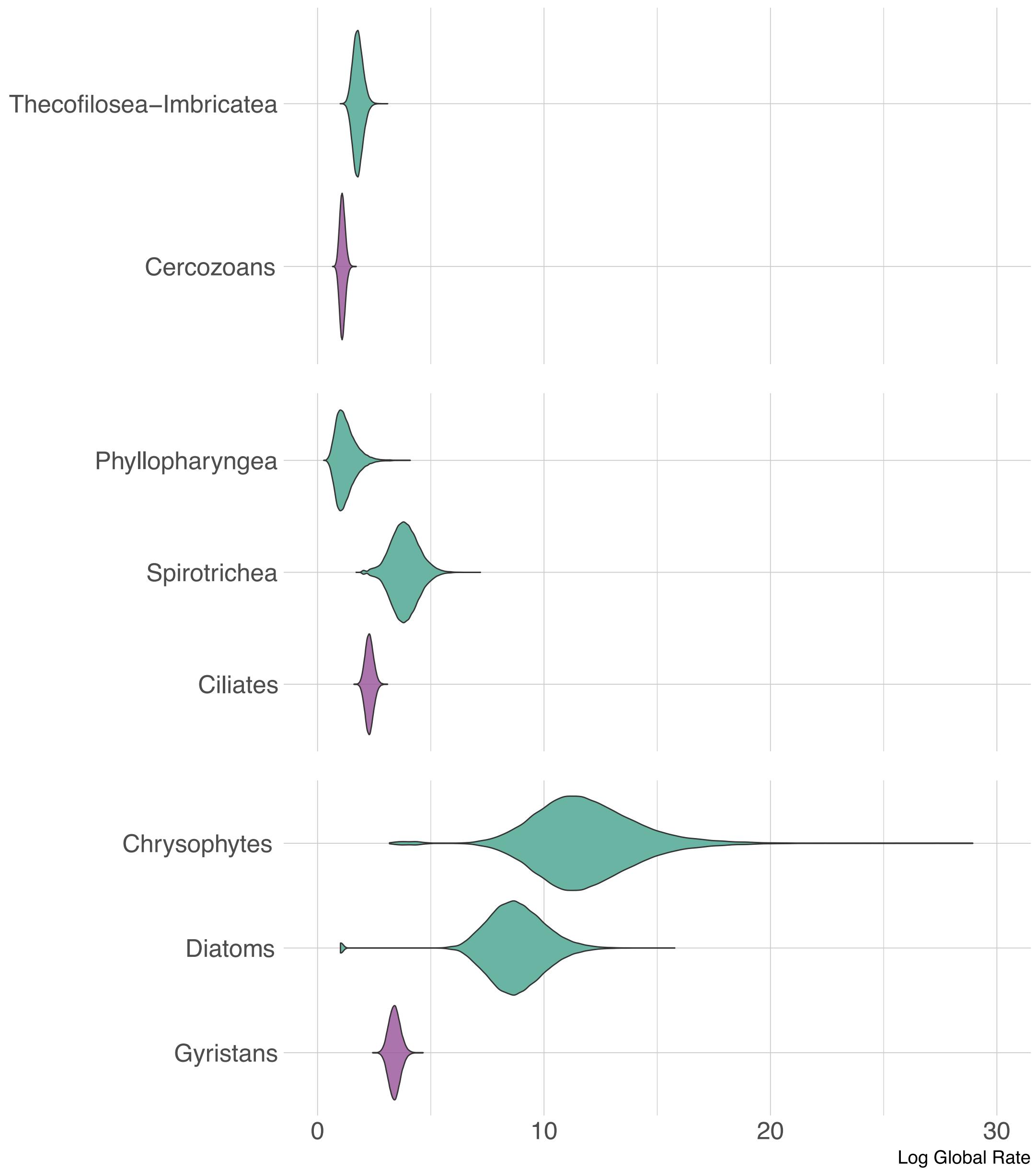

**Supplementary Figure 11.** Posterior probability distributions of the global habitat evolution rates of selected eukaryotic classes. In purple, are the global transition rates of three major eukaryotic lineages (Cercozoans, Ciliates, Gyristans) as shown in Figure 2, and in green, are the global transition rates for selected clades within these lineages. From this analysis, we can see that Thecofilosea+Imbricatea tend to transition across the salt-barrier faster than Cercozoans on the whole. Spirotrichea have higher transition rates than Ciliates on average, and Chrysophtes (golden algae) and Diatoms seem to have the highest transition rates across protists.

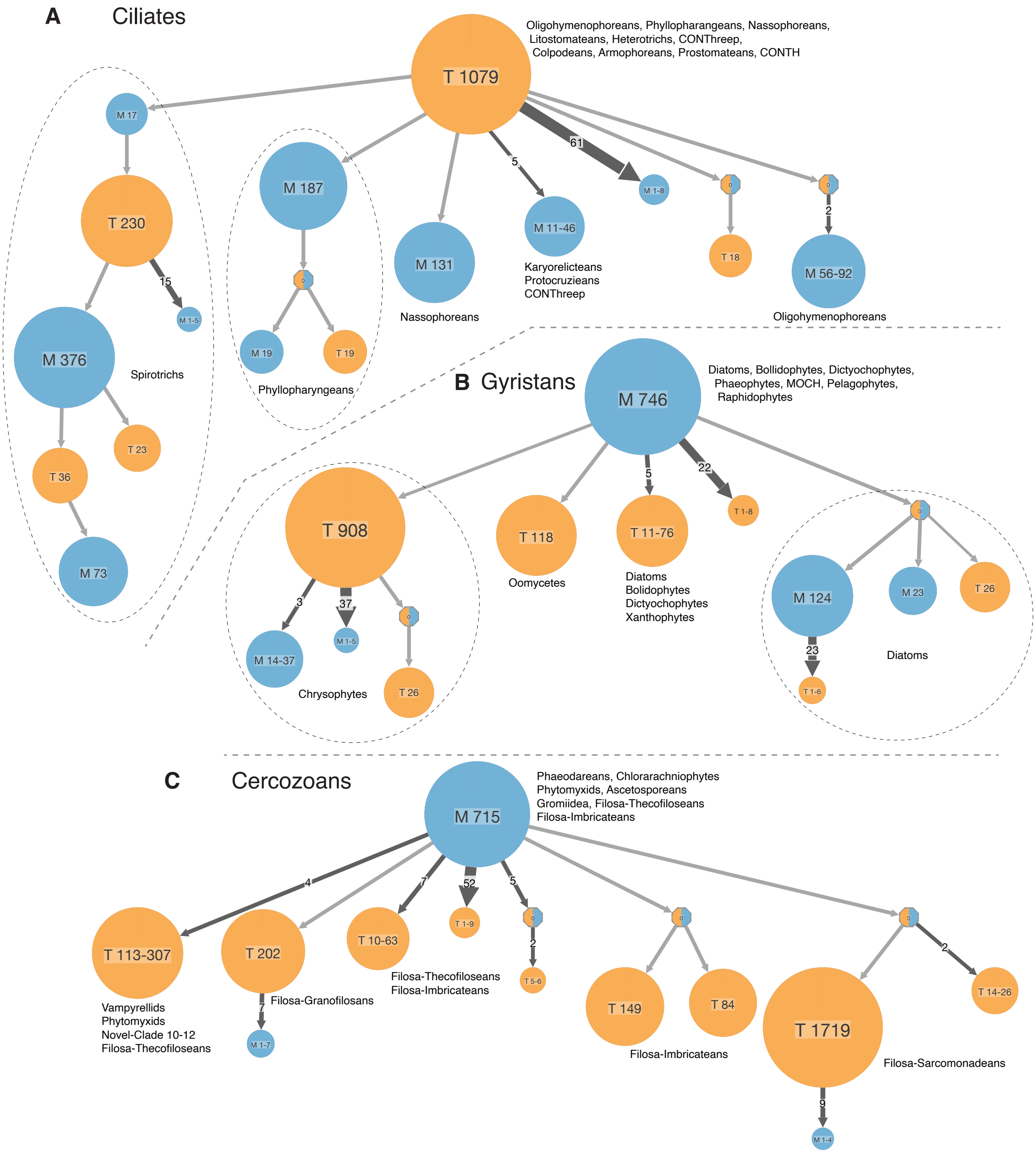

**Supplementary Figure 12.** Simplified possible ancestral scenarios depicted for (**A**) Ciliates, (**B**) Gyristans, and (**C**) Cercozoans as generated by PASTML using marginal posterior probability approximation (MPPA) with an F-81 like model. For this compressed visualisation, parts of the tree with no state changes are collapsed into circles, with the number in each circle indicating the number of taxa. Orange circles depict terrestrial clades, and blue circles depict marine clades. We display the eukaryotic classes represented in each circle, for instance, the first orange circle in ciliates indicates a terrestrial origin, with 1079 taxa representing oligohymenophoreans, nassophoreans, colpodeans and many other ciliate classes. Thin grey lines represent a single transition event. For example, Panel **A** shows a single terrestrial to marine transition leading to spirotrichs. Thick lines represent multiple transition events with the associated number indicating the number of such events. For example, within spirotrichs (Panel **A**), there were 15 transition events leading to marine clades of sizes one to five taxa. For the sake of simplification, PASTML displays only major transition events and hides minor ones (further from the root). Therefore certain transition events are not depicted, such as transitions from marine to terrestrial environments in chrysophytes, and such as transitions from terrestrial to marine environments in vampyrellids.

This figure depicts the high number of serial transitions in spirotrich ciliates for which we can observe repeated re-colonizations of marine and terrestrial habitats. Also depicted are a high number of independent transitions within diatoms, chryosphytes (in Panel **B**), and thecofiloseans and imbricateans (in Panel **C**) in accordance with Supplementary Figure 11.

This figure also shows transition events which led to the establishment of several key-deep branching lineages. This pattern is best exemplified by cercozoans, where one putative transition from marine to terrestrial environments led to the diversification of vampyrellids, a group of predatory amoebas which hunt algae (Panel **C**); and another putative transition in the same direction led to the largely uncharacterized Novel-10-12 clade[7] which is commonly found in freshwaters.

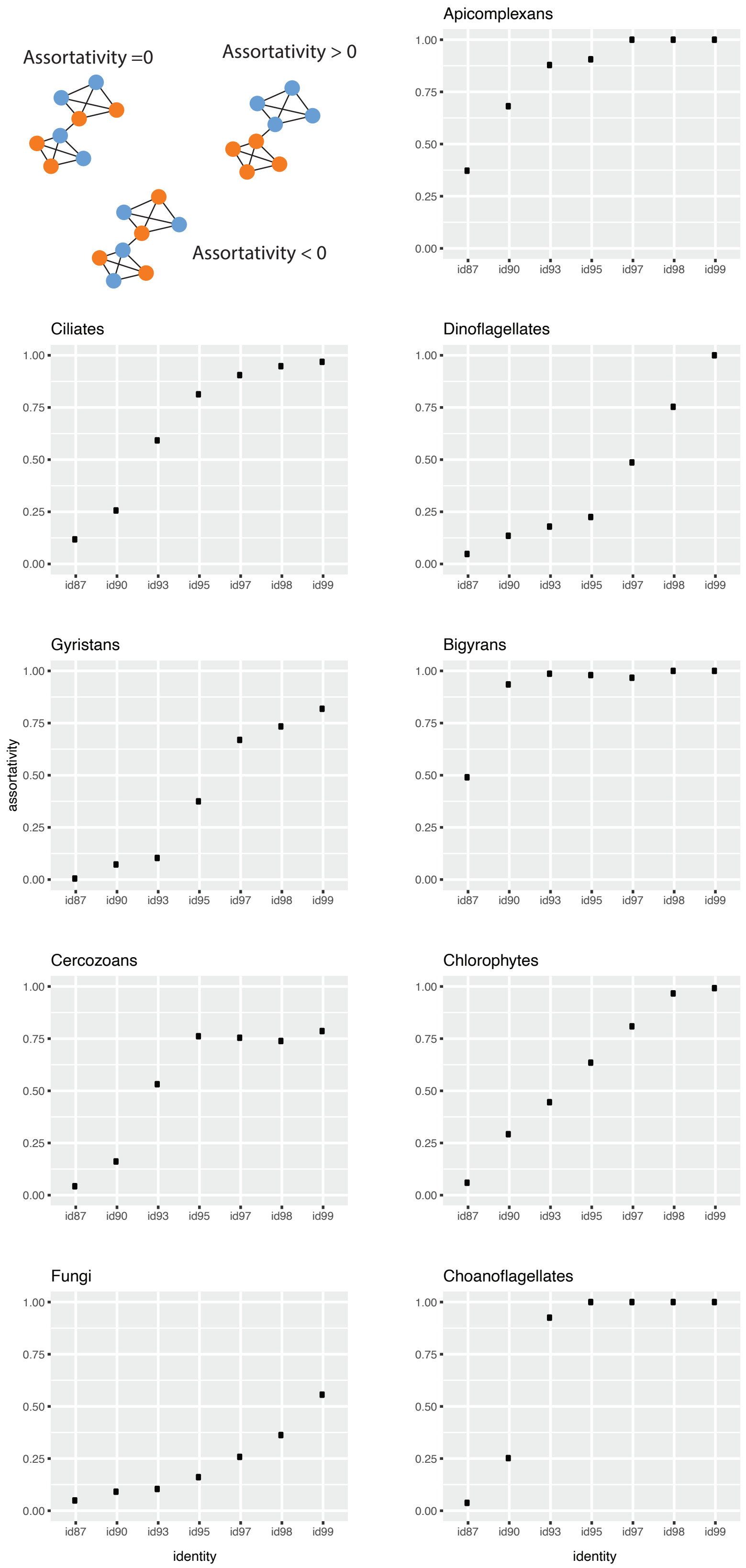

**Supplementary Figure 13.** Sequence similarity network analyses on PacBio sequences. 18S sequences were clustered at different similarity thresholds to produce networks. A simple representation is shown in the top right panel, where each node represents a sequence, and each edge represents the connection between nodes. The assortativity of marine and terrestrial sequences was measured, which measures whether nodes with the same attribute tend to connect with each other more than random or not. When assortatvity = 0, that indicates, in this case, that marine and terrestrial nodes connect to each other randomly. When assortativity tends towards 1, that means that marine and terrestrial nodes tend to preferentially connect to nodes from the same habitat. With these analyses, we can see that Fungi have the lowest assortativity values. This can also be taken to mean that fungal transitions have occurred very recently, so that ribosomal sequences have not had enough time to evolve since the transition.

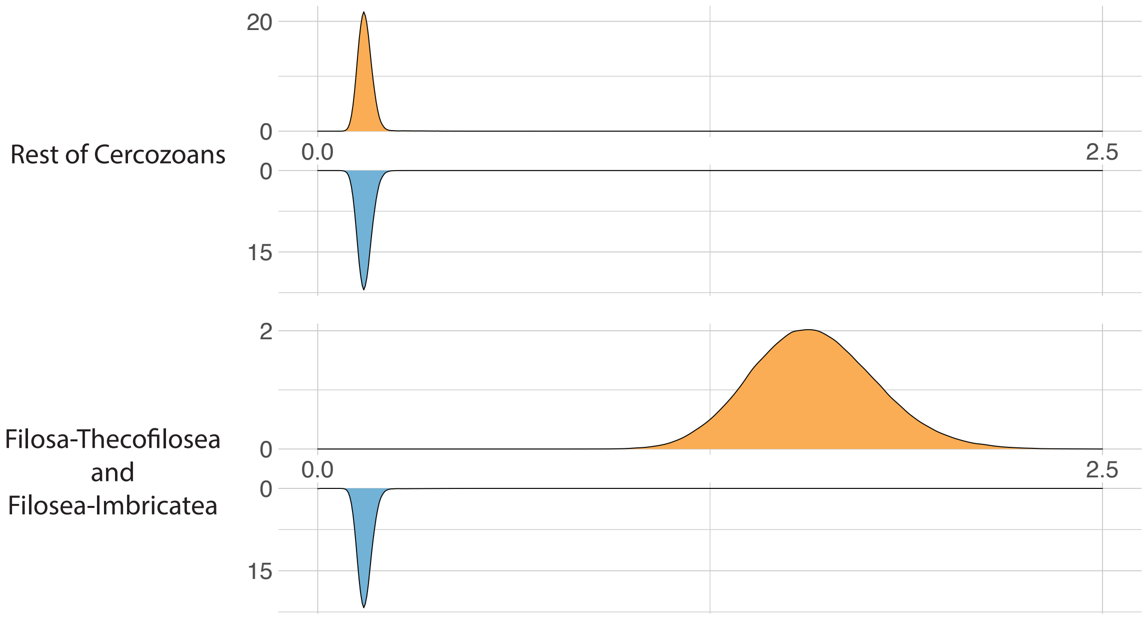

**Supplementary Figure 14.** Habitat transition rates vary within Cercozoans. Here, we used a heterogenous model of habitat evolution such that qMT and qTM were estimated separately for Filosa-Thecofilosea + Filosa-Imbricatea, and the rest of Cercozoans. Filosa-Thecofilosea + Filosa-Imbricatea was selected based on visual inspection of the Cercozoan phylogenies (where terrestrial and marine lineages were more interspersed and not as distinct).

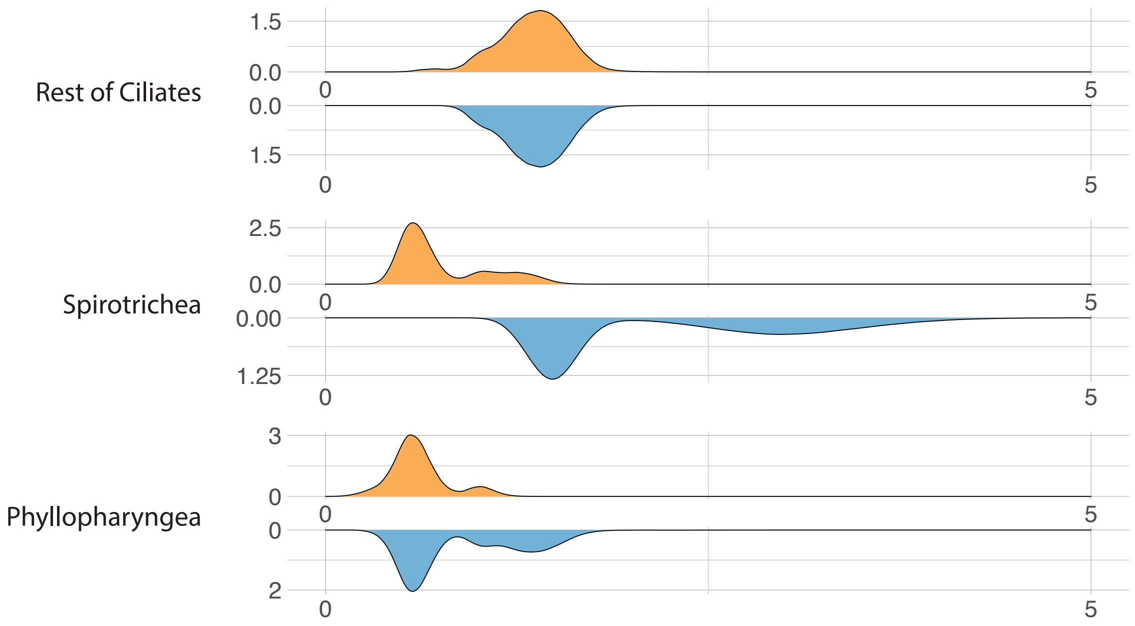

**Supplementary Figure 15.** Habitat transition rates vary within Ciliates. Here, we used a heterogenous model of habitat evolution such that qMT and qTM were estimated separately for Spirotrichea, Phyllopharyngea and the rest of Ciliates. Spirotrichea and Phyllopharyngea were selected as they showed multiple putative serial colonization events (see Figure 5).

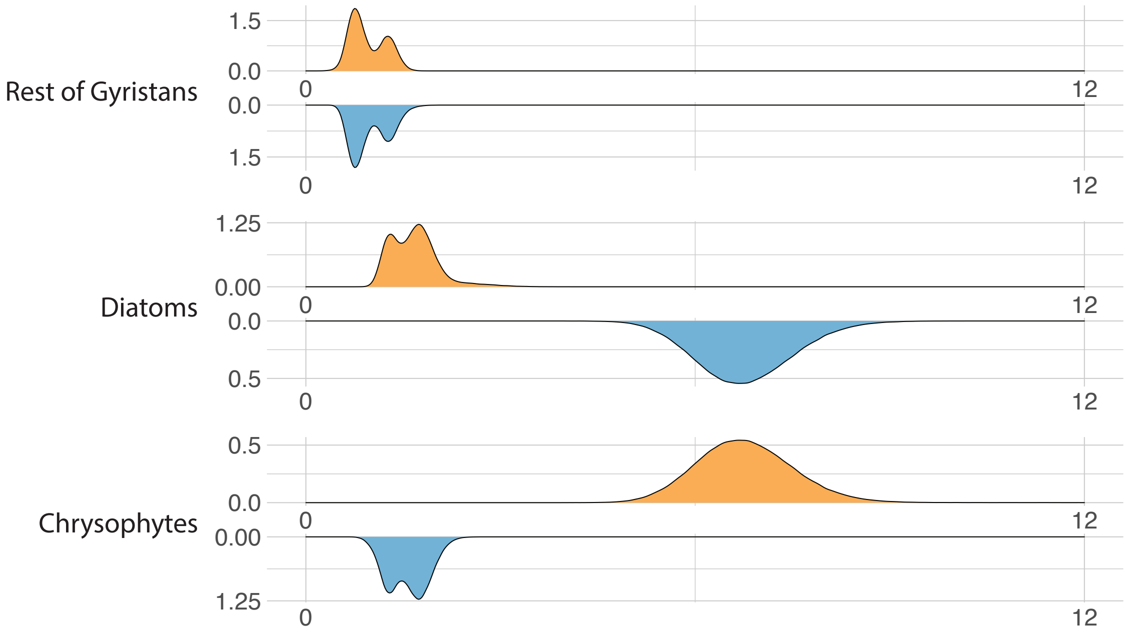

**Supplementary Figure 16.** Habitat transition rates vary within Gyristans. Here, we used a heterogenous model of habitat evolution such that qMT and qTM were estimated separately for Diatoms, Chrysophytes and the rest of Gyristans. Diatoms and Chrysophytes seemed to have less distinct marine and terrestrial lineages based on visual inspection of the Gyristan phylogenies, hence we chose to characterize these clades separately.

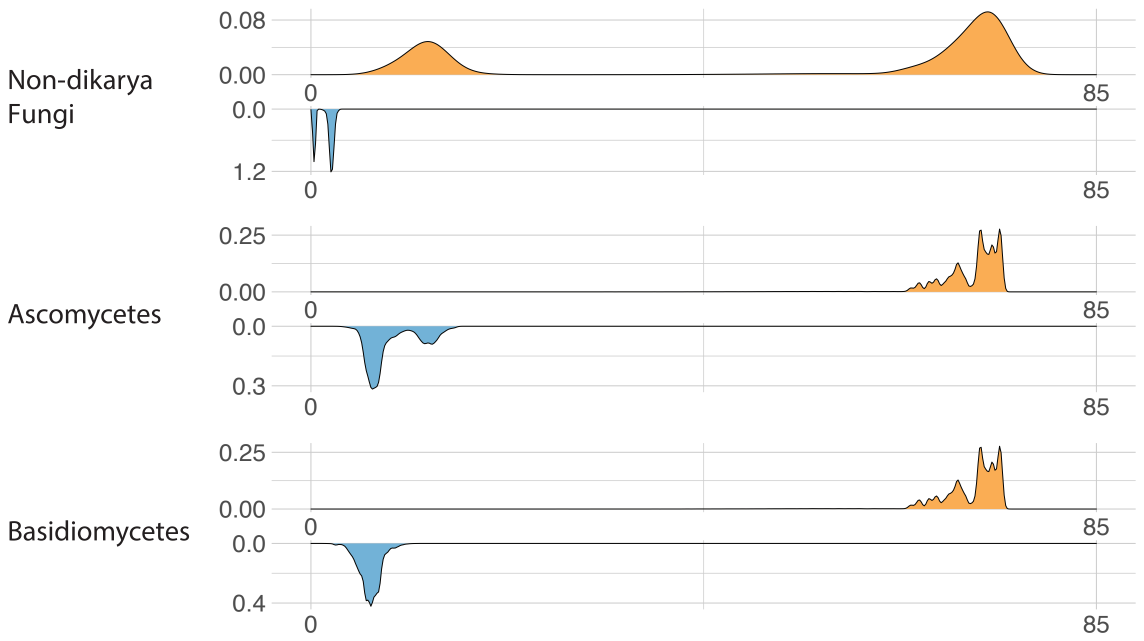

**Supplementary Figure 17.** Habitat transition rates vary within Fungi. Here, we used a heterogenous model of habitat evolution such that qMT and qTM were estimated separately for Ascomycetes, Basidiomycetes, and non-Dikarya Fungi. Visual inspection of fungal phylogenies indicated more transition events in Dikarya as compared to non-Dikarya.

**Supplementary Note 1. Estimate of ancestral habitats of eukaryotic groups with insufficient PacBio data.**

**Collodictyonids**. *Collodictyon triciliatum* was first isolated and described from freshwater habitats[8–10]. An environmental survey using group specific primers revealed a global distribution that was limited to freshwater environments[11]. No collodictyonid has been found in marine environments, therefore a terrestrial origin seems more likely.

**Metamonads.** Mantamonads are gliding flagellates that have thus far only been found in benthic marine habitats[12,13].

**Pluriformeans.** This group currently consists of two genera: *Corallochytrium* which is a marine organism associated with corals, and *Syssomonas* which is found in freshwater. The ancestral habitat of this group is thus ambiguous[14].

**Ancoracystids.** The only described organism from this group, *Ancoracysta twista*, was isolated from the surface of a marine coral[15].

**Glaucophytes.** Almost all glaucophyte algae inhabit freshwater environments, making a terrestrial origin more likely[16–18].

**Rhodophytes.** While several rhodophytes are found in freshwater lakes and even soils, the vast majority of rhodophytes are marine, and ancestral state reconstruction studies indicate a marine origin[17,19].

**Rhodelphids.** Two species have been described from this group thus far: *Rhodelphis marinus* which was obtained from marine coral sand, and *Rhodelphis limneticus* which was obtained from a freshwater lake[20]. The ancestral habitat of the group is therefore ambiguous.

**Supplementary Table 1.** Collection details of the 21 samples sequenced with PacBio Sequel II for this study. All raw sequencing data can be accessed at ENA under accession PRJEB45931 (this study) and PRJEB25197 (sequenced in [21]). SITES = Swedish Infrastructure for Ecosystem Science. SMHI = Swedish Meteorological and Hydrological Institute.

| No. | Sample code | Sample type, level 1 | Sample type, level 2 | Sampling date (yyyy-mm-dd) | Sample collected by | No. of sites pooled | Country/Region | Sampling Location | Comments | Size fraction | Lat | Long | Depth | Reference for samples | Sequencing instrument | No. of demultiplexed reads | No. of processed reads | No. of OTUs |
|---|---|---|---|---|---|---|---|---|---|---|---|---|---|---|---|---|---|---|
| 1 | Erken_fw | terrestrial | freshwater | 2019-10-09 | SITES | 1 | Sweden | Lake Erken, Erken research station | Hemi-boreal, agricultural landscape | 200-0.2 µm | 59.83 | 18.63 | 0-20 m | This study | Sequel II | 564,504 | 320,773 | 768 |
| 2 | Skogaryd_fw | terrestrial | freshwater | 2019-10-17 | SITES | 1 | Sweden | Lake Ersjön, Skogaryd research station | Hemi-boreal, forest landscape | 200-0.2 µm | 58.37 | 12.16 | 0.5, 2, 4 m | This study | Sequel II | 699,716 | 422,836 | 1688 |
| 3 | Svartberget_fw | terrestrial | freshwater | 2019-11-05 | SITES | 1 | Sweden | Lake Stortjärn, Svartberget research station | Boreal, forest landscape | 200-0.2 µm | 64.24 | 19.76 | 0.5, 3.5 m | This study | Sequel II | 627,565 | 373,152 | 496 |
| 4 | Permafrost_fw | terrestrial | freshwater | 2014-08 | S. Peura, K. Einarsdottir, M. Wauthy | 10 | Canada | Nunavik, Quebec | Multiple permafrost thaw ponds | > 0.2 µm | 55.22 | -77.69 | 0-1.7 m | [22] | Sequel II | 456,683 | 256,770 | 1324 |
| 5 | Erken_sed_fw | terrestrial | freshwater | 2019-10-09 | SITES | 4 | Sweden | Lake Erken, Erken research station | Sediment core of depth 0-5 cm | No size selection | 59.83 | 18.63 | 0-5 cm | This study | Sequel II | 519,814 | 337,656 | 1391 |
| 6 | Skogaryd_soil | terrestrial | soil | 2019-10-17 | SITES | 4 | Sweden | Skogaryd Mire, Skogaryd research station | Composite sample from wet and dry areas | No size selection | 58.37 | 12.16 | top soil | This study | Sequel II | 301,806 | 90,907 | 593 |
| 7 | Svartberget_soil | terrestrial | soil | 2019-11-05 | SITES | 3 | Sweden | Kallkäls Mire, Svartberget research station |  | No size selection | 64.24 | 19.76 | top soil | This study | Sequel II | 241,901 | 133,266 | 763 |
| 8 | PuertoRico_soil | terrestrial | soil | 2013-12 | H. Urbina | 6 | Puerto Rico | El Yunque, Puerto Rico | Montane wet forest | No size selection | 18.29 | -65.78 | top soil | [23] | Sequel II | 770,855 | 384,698 | 441 |
| 9 | Sweden_soil | terrestrial | soil | 2013-10-15 | H. Urbina | 6 | Sweden | Ivantjärnsheden field station, Jädraås | Pine forest | No size selection | 60.49 | 16.3 | top soil | [24] | Sequel II | 950,141 | 519,140 | 1184 |
| 10 | agricultural_soil | terrestrial | soil | 2008-03-04 | P. Gosling, G. Bending | 14 | UK | Set aside arable sites, UK | Sequenced in previous study | No size selection | 51.21 | 1.23 | top soil | [21] | Sequel | 36,339 | 10,755 | 595 |
| 11 | rhizosphere_soil | terrestrial | soil | 2015-03 | S. Hilton | 33 | UK | 25 commercial farms in the UK | Sequenced on Sequel I | No size selection | 52.2 | -0.81 | top soil | [21] | Sequel | 28,347 | 9,013 | 222 |
| 12 | Tibet_soil | terrestrial | soil | 06/08-2011 | S. Geisen, J. Zhang | 9 | Tibet | Mix of alpine meadows, shrubs, and forest | Sequenced in previous study | No size selection | 29.36 | 94.43 | top soil | [21] | Sequel | 48,676 | 4,315 | 183 |
| 13 | WC_sur | marine | euphotic | 2019-08-20 | SMHI | 1 | North Sea | Station Anholt E |  | 200-0.2 µm | 56.66 | 12.11 | 5 m | This study | Sequel II | 1,543,739 | 964,198 | 639 |
| 14 | Ms1_DCM | marine | euphotic | 2011-02-17 | Malaspina team | 2 | Indian Ocean | Malaspina station 49 | DCM layer | 3-0.2 µm | -33.9 | 37 | 85 m | [25] | Sequel II | 273,073 | 167,358 | 1126 |
| 15 | Ms2_sur | marine | euphotic | 2011-03-25 | Malaspina team | 2 | Indian Ocean | Malaspina station 76 | Surface | 3-0.2 µm | -40.6 | 142.5 | 3 m | [25] | Sequel II | 149,790 | 93,932 | 377 |
| 16 | Ms2_DCM | marine | euphotic | 2011-03-25 | Malaspina team | 2 | Indian Ocean | Malaspina station 76 | DCM layer | 20-0.2 µm | -40.6 | 142.5 | 70 m | [25] | Sequel II | 511,880 | 272,269 | 754 |
| 17 | Ms1_mes | marine | aphotic | 2011-02-17 | Malaspina team | 2 | Indian Ocean | Malaspina station 49 | Mesopelagic | 3-0.2 µm | -33.9 | 37 | 800 m | [25] | Sequel II | 642,804 | 243,523 | 540 |
| 18 | Ms2_mes | marine | aphotic | 2011-03-25 | Malaspina team | 2 | Indian Ocean | Malaspina station 76 | Mesopelagic | 20-0.2 µm | -40.6 | 142.5 | 275 m | [25] | Sequel II | 656,429 | 415,020 | 637 |
| 19 | Ms1_bat | marine | aphotic | 2011-02-17 | Malaspina team | 2 | Indian Ocean | Malaspina station 49 | Bathypelagic | 20-0.2 µm | -33.9 | 37 | 1200 m | [25] | Sequel II | 761,055 | 422,088 | 1554 |
| 20 | Ms2_bat | marine | aphotic | 2011-03-25 | Malaspina team | 2 | Indian Ocean | Malaspina station 76 | Bathypelagic | 20-0.2 µm | -40.6 | 142.5 | 2800-3300 m | [25] | Sequel II | 775,615 | 483,155 | 1196 |
| 21 | MT | marine | aphotic | 2016-06 | H. Jing, pilots of Jiao Long Hao | 1 | Pacific Ocean | Mariana Trench | Bathypelagic | 3-0.2 µm | 11.18 | 141.98 | 5900 m | [26] | Sequel II | 162,569 | 84,921 | 99 |

**Supplementary Table 2.** The first five ranks of the PR2-transitions database[27] (upto the class rank; see Materials and Methods for details on how the PR2 database was adapted for this study), and whether corresponding ribosomal DNA sequences were recovered by PacBio sequencing.

| Domain | Supergroup | Division | Subdivision | Class | Sequenced by PacBio |
|---|---|---|---|---|---|
| Eukaryota | Amoebozoa | Amoebozoa_X | Amoebozoa_XX | Amoebozoa_XXX | Yes |
| Eukaryota | Amoebozoa | Amoebozoa_X | Amoebozoa_XX | Lobosa-G1 | Yes |
| Eukaryota | Amoebozoa | Discosea | Discosea_X | Centramoebia | Yes |
| Eukaryota | Amoebozoa | Discosea | Discosea_X | Flabellinia | Yes |
| Eukaryota | Amoebozoa | Discosea | Discosea_X | Stygamoebida | Yes |
| Eukaryota | Amoebozoa | Evosea | Evosea_X | Archamoebea | Yes |
| Eukaryota | Amoebozoa | Evosea | Evosea_X | Eumycetozoa | Yes |
| Eukaryota | Amoebozoa | Evosea | Evosea_X | Variosea | Yes |
| Eukaryota | Amoebozoa | Tubulinea | Tubulinea_X | Corycida | - |
| Eukaryota | Amoebozoa | Tubulinea | Tubulinea_X | Echinamoebida | Yes |
| Eukaryota | Amoebozoa | Tubulinea | Tubulinea_X | Elardia | Yes |
| Eukaryota | Amoebozoa | Tubulinea | Tubulinea_X | Tubulinea_XX | Yes |
| Eukaryota | Archaeplastida | Chlorophyta | Chlorophyta_X | Chlorodendrophyceae | Yes |
| Eukaryota | Archaeplastida | Chlorophyta | Chlorophyta_X | Chlorophyceae | Yes |
| Eukaryota | Archaeplastida | Chlorophyta | Chlorophyta_X | Chlorophyta_XX | Yes |
| Eukaryota | Archaeplastida | Chlorophyta | Chlorophyta_X | Chloropicophyceae | Yes |
| Eukaryota | Archaeplastida | Chlorophyta | Chlorophyta_X | Mamiellophyceae | Yes |
| Eukaryota | Archaeplastida | Chlorophyta | Chlorophyta_X | Nephroselmidophyceae | - |
| Eukaryota | Archaeplastida | Chlorophyta | Chlorophyta_X | Palmophyllophyceae | Yes |
| Eukaryota | Archaeplastida | Chlorophyta | Chlorophyta_X | Pedinophyceae | Yes |
| Eukaryota | Archaeplastida | Chlorophyta | Chlorophyta_X | Picocystophyceae | - |
| Eukaryota | Archaeplastida | Chlorophyta | Chlorophyta_X | Prasino-Clade-9 | Yes |
| Eukaryota | Archaeplastida | Chlorophyta | Chlorophyta_X | Prasino-Clade-VIII | - |
| Eukaryota | Archaeplastida | Chlorophyta | Chlorophyta_X | Prasino-Clade-V | Yes |
| Eukaryota | Archaeplastida | Chlorophyta | Chlorophyta_X | Pyramimonadophyceae | Yes |
| Eukaryota | Archaeplastida | Chlorophyta | Chlorophyta_X | Trebouxiophyceae | Yes |
| Eukaryota | Archaeplastida | Chlorophyta | Chlorophyta_X | Ulvophyceae | Yes |
| Eukaryota | Archaeplastida | Glaucophyta | Glaucophyta_X | Glaucocystophyceae | Yes |
| Eukaryota | Archaeplastida | Rhodelphea | Rhodelphea_X | Rhodelphida | Yes |
| Eukaryota | Archaeplastida | Rhodophyta | Rhodophyta_X | Bangiophyceae | Yes |
| Eukaryota | Archaeplastida | Rhodophyta | Rhodophyta_X | Compsopogonophyceae | - |

| Domain | Supergroup | Division | Subdivision | Class | Sequenced by PacBio |
|---|---|---|---|---|---|
| Eukaryota | Archaeplastida | Rhodophyta | Rhodophyta_X | Florideophyceae | - |
| Eukaryota | Archaeplastida | Rhodophyta | Rhodophyta_X | Porphyridiophyceae | - |
| Eukaryota | Archaeplastida | Rhodophyta | Rhodophyta_X | Rhodellophyceae | - |
| Eukaryota | Archaeplastida | Rhodophyta | Rhodophyta_X | Rhodophyta_XX | - |
| Eukaryota | Archaeplastida | Rhodophyta | Rhodophyta_X | Stylonematophyceae | - |
| Eukaryota | Archaeplastida | Streptophyta | Streptophyta_X | Charophyceae | - |
| Eukaryota | Archaeplastida | Streptophyta | Streptophyta_X | Coleochaetophyceae | - |
| Eukaryota | Archaeplastida | Streptophyta | Streptophyta_X | Embryophyceae | Yes |
| Eukaryota | Archaeplastida | Streptophyta | Streptophyta_X | Klebsormidiophyceae | Yes |
| Eukaryota | Archaeplastida | Streptophyta | Streptophyta_X | Mesostigmatophyceae | - |
| Eukaryota | Archaeplastida | Streptophyta | Streptophyta_X | Streptophyta_XX | Yes |
| Eukaryota | Archaeplastida | Streptophyta | Streptophyta_X | Zygnemophyceae | Yes |
| Eukaryota | CRuMs | Collodictyonidae | Collodictyonidae_X | Collodictyonidae_XX | - |
| Eukaryota | CRuMs | Mantamonadidea | Mantamonadidea_X | Mantamonadida | - |
| Eukaryota | CRuMs | Rigifilida | Rigifilida_X | Rigifilida_XX | Yes |
| Eukaryota | Cryptista | Cryptista_X | Cryptista_XX | Cryptista_XXX | - |
| Eukaryota | Cryptista | Cryptophyta | Cryptophyta_X | Cryptophyceae | Yes |
| Eukaryota | Cryptista | Kathablepharidacea | Kathablepharida | Kathablepharidea | Yes |
| Eukaryota | Eukaryota_X | Ancoracystida | Ancoracystida_X | Ancoracystida_XX | - |
| Eukaryota | Eukaryota_X | Ancyromonadida | Ancyromonadida_X | Ancyromonadida_XX | Yes |
| Eukaryota | Eukaryota_X | Hemimastigophora | Hemimastigophora_X | Hemimastigophora_XX | Yes |
| Eukaryota | Eukaryota_X | Picozoa | Picozoa_X | Picozoa_XX | Yes |
| Eukaryota | Excavata | Discoba | Discoba_X | Euglenozoa | Yes |
| Eukaryota | Excavata | Discoba | Discoba_X | Heterolobosea | Yes |
| Eukaryota | Excavata | Discoba | Discoba_X | Jakobida | Yes |
| Eukaryota | Excavata | Discoba | Discoba_X | Tsukubamonadidae | - |
| Eukaryota | Excavata | Malawimonadidae | Malawimonadidae_X | Malawimonadidae_XX | Yes |
| Eukaryota | Excavata | Metamonada | Metamonada_X | Fornicata | Yes |
| Eukaryota | Excavata | Metamonada | Metamonada_X | Parabasalia | - |
| Eukaryota | Excavata | Metamonada | Metamonada_X | Preaxostyla | Yes |
| Eukaryota | Haptista | Centroplasthelida | Centroplasthelida_X | Centroplasthelida_XX | Yes |
| Eukaryota | Haptista | Centroplasthelida | Centroplasthelida_X | Panacanthocystida | Yes |
| Eukaryota | Haptista | Centroplasthelida | Centroplasthelida_X | Pterocystida | Yes |
| Eukaryota | Haptista | Haptophyta | Haptophyta_X | Haptophyta_Clade_HAP1 | - |

| Domain | Supergroup | Division | Subdivision | Class | Sequenced by PacBio |
|---|---|---|---|---|---|
| Eukaryota | Haptista | Haptophyta | Haptophyta_X | Haptophyta_Clade_HAP2 | Yes |
| Eukaryota | Haptista | Haptophyta | Haptophyta_X | Haptophyta_Clade_HAP3 | Yes |
| Eukaryota | Haptista | Haptophyta | Haptophyta_X | Haptophyta_Clade_HAP4 | Yes |
| Eukaryota | Haptista | Haptophyta | Haptophyta_X | Haptophyta_Clade_HAP5 | - |
| Eukaryota | Haptista | Haptophyta | Haptophyta_X | Haptophyta_XX | Yes |
| Eukaryota | Haptista | Haptophyta | Haptophyta_X | Pavlovophyceae | Yes |
| Eukaryota | Haptista | Haptophyta | Haptophyta_X | Prymnesiophyceae | Yes |
| Eukaryota | Obazoa | Apusomonada | Apusomonada_X | Apusomonadidae | Yes |
| Eukaryota | Obazoa | Breviatea | Breviatea_X | Breviatea_XX | Yes |
| Eukaryota | Obazoa | Breviatea | Breviatea_X | NAMAKO-1-lineage | - |
| Eukaryota | Obazoa | Breviatea | Breviatea_X | YS16Ec34-lineage | - |
| Eukaryota | Obazoa | Opisthokonta | Choanoflagellata | Choanoflagellata_X | Yes |
| Eukaryota | Obazoa | Opisthokonta | Filasterea | Filasterea_X | Yes |
| Eukaryota | Obazoa | Opisthokonta | Fungi | Ascomycota | Yes |
| Eukaryota | Obazoa | Opisthokonta | Fungi | Basidiomycota | Yes |
| Eukaryota | Obazoa | Opisthokonta | Fungi | Blastocladiomycota | Yes |
| Eukaryota | Obazoa | Opisthokonta | Fungi | Chytridiomycota | Yes |
| Eukaryota | Obazoa | Opisthokonta | Fungi | Fungi_X | Yes |
| Eukaryota | Obazoa | Opisthokonta | Fungi | Monoblepharidomycetes | Yes |
| Eukaryota | Obazoa | Opisthokonta | Fungi | Mucoromycota | Yes |
| Eukaryota | Obazoa | Opisthokonta | Fungi | Neocallimastigaceae | - |
| Eukaryota | Obazoa | Opisthokonta | Fungi | Olpidium_class | - |
| Eukaryota | Obazoa | Opisthokonta | Fungi | Opisthosporidia | Yes |
| Eukaryota | Obazoa | Opisthokonta | Fungi | Zoopagomycota | Yes |
| Eukaryota | Obazoa | Opisthokonta | Ichthyosporea | Dermocystida | Yes |
| Eukaryota | Obazoa | Opisthokonta | Ichthyosporea | Ichthyophonida | Yes |
| Eukaryota | Obazoa | Opisthokonta | Ichthyosporea | Ichthyosphonida | Yes |
| Eukaryota | Obazoa | Opisthokonta | Metazoa | Acanthocephala | Yes |
| Eukaryota | Obazoa | Opisthokonta | Metazoa | Annelida | Yes |
| Eukaryota | Obazoa | Opisthokonta | Metazoa | Arthropoda | Yes |
| Eukaryota | Obazoa | Opisthokonta | Metazoa | Arthropoda | Yes |
| Eukaryota | Obazoa | Opisthokonta | Metazoa | Arthropoda | Yes |
| Eukaryota | Obazoa | Opisthokonta | Metazoa | Arthropoda | Yes |
| Eukaryota | Obazoa | Opisthokonta | Metazoa | Arthropoda | Yes |

| Domain | Supergroup | Division | Subdivision | Class | Sequenced by PacBio |
|---|---|---|---|---|---|
| Eukaryota | Obazoa | Opisthokonta | Metazoa | Brachiopoda | Yes |
| Eukaryota | Obazoa | Opisthokonta | Metazoa | Bryozoa | Yes |
| Eukaryota | Obazoa | Opisthokonta | Metazoa | Cephalochordata | - |
| Eukaryota | Obazoa | Opisthokonta | Metazoa | Chaetognatha | Yes |
| Eukaryota | Obazoa | Opisthokonta | Metazoa | Cnidaria | Yes |
| Eukaryota | Obazoa | Opisthokonta | Metazoa | Craniata | - |
| Eukaryota | Obazoa | Opisthokonta | Metazoa | Ctenophora | Yes |
| Eukaryota | Obazoa | Opisthokonta | Metazoa | Cycliophora | - |
| Eukaryota | Obazoa | Opisthokonta | Metazoa | Echinodermata | Yes |
| Eukaryota | Obazoa | Opisthokonta | Metazoa | Entoprocta | - |
| Eukaryota | Obazoa | Opisthokonta | Metazoa | Gastrotricha | Yes |
| Eukaryota | Obazoa | Opisthokonta | Metazoa | Gnathostomulida | - |
| Eukaryota | Obazoa | Opisthokonta | Metazoa | Hemichordata | - |
| Eukaryota | Obazoa | Opisthokonta | Metazoa | Kinorhyncha | - |
| Eukaryota | Obazoa | Opisthokonta | Metazoa | Loricifera | - |
| Eukaryota | Obazoa | Opisthokonta | Metazoa | Mesozoa | - |
| Eukaryota | Obazoa | Opisthokonta | Metazoa | Metazoa_X | - |
| Eukaryota | Obazoa | Opisthokonta | Metazoa | Micrognathozoa | - |
| Eukaryota | Obazoa | Opisthokonta | Metazoa | Mollusca | Yes |
| Eukaryota | Obazoa | Opisthokonta | Metazoa | Myxozoa | Yes |
| Eukaryota | Obazoa | Opisthokonta | Metazoa | Myzostomida | - |
| Eukaryota | Obazoa | Opisthokonta | Metazoa | Nematoda | Yes |
| Eukaryota | Obazoa | Opisthokonta | Metazoa | Nematomorpha | Yes |
| Eukaryota | Obazoa | Opisthokonta | Metazoa | Nemertea | - |
| Eukaryota | Obazoa | Opisthokonta | Metazoa | Onychophora | - |
| Eukaryota | Obazoa | Opisthokonta | Metazoa | Placozoa | - |
| Eukaryota | Obazoa | Opisthokonta | Metazoa | Platyhelminthes | Yes |
| Eukaryota | Obazoa | Opisthokonta | Metazoa | Porifera | - |
| Eukaryota | Obazoa | Opisthokonta | Metazoa | Priapulida | - |
| Eukaryota | Obazoa | Opisthokonta | Metazoa | Rotifera | Yes |
| Eukaryota | Obazoa | Opisthokonta | Metazoa | Tardigrada | Yes |
| Eukaryota | Obazoa | Opisthokonta | Metazoa | Urochordata | Yes |
| Eukaryota | Obazoa | Opisthokonta | Metazoa | Xenoturbellida | - |
| Eukaryota | Obazoa | Opisthokonta | Opisthokonta_X | Opisthokonta_XX | Yes |

| Domain | Supergroup | Division | Subdivision | Class | Sequenced by PacBio |
|---|---|---|---|---|---|
| Eukaryota | Obazoa | Opisthokonta | Pluriformea | Corallochytrea | - |
| Eukaryota | Obazoa | Opisthokonta | Pluriformea | Pluriformea_X | Yes |
| Eukaryota | Obazoa | Opisthokonta | Rotosphaerida | Fonticulea | - |
| Eukaryota | Obazoa | Opisthokonta | Rotosphaerida | Nucleariidea | Yes |
| Eukaryota | TSAR | Alveolata | Alveolata_X | Alveolata_XX | - |
| Eukaryota | TSAR | Alveolata | Apicomplexa | Apicomplexa_X | Yes |
| Eukaryota | TSAR | Alveolata | Apicomplexa | Coccidiomorphea | Yes |
| Eukaryota | TSAR | Alveolata | Apicomplexa | Colpodellidea | Yes |
| Eukaryota | TSAR | Alveolata | Apicomplexa | Gregarinomorphea | Yes |
| Eukaryota | TSAR | Alveolata | Ciliophora | Armophorea | Yes |
| Eukaryota | TSAR | Alveolata | Ciliophora | Cariacotrichea | - |
| Eukaryota | TSAR | Alveolata | Ciliophora | Ciliophora_X | Yes |
| Eukaryota | TSAR | Alveolata | Ciliophora | Colpodea | Yes |
| Eukaryota | TSAR | Alveolata | Ciliophora | CONTH_1 | - |
| Eukaryota | TSAR | Alveolata | Ciliophora | CONTH_2 | Yes |
| Eukaryota | TSAR | Alveolata | Ciliophora | CONTH_3 | - |
| Eukaryota | TSAR | Alveolata | Ciliophora | CONTH_4 | Yes |
| Eukaryota | TSAR | Alveolata | Ciliophora | CONTH_5 | Yes |
| Eukaryota | TSAR | Alveolata | Ciliophora | CONTH_6 | - |
| Eukaryota | TSAR | Alveolata | Ciliophora | CONTH_7 | Yes |
| Eukaryota | TSAR | Alveolata | Ciliophora | CONTH_8 | - |
| Eukaryota | TSAR | Alveolata | Ciliophora | CONThreeP | Yes |
| Eukaryota | TSAR | Alveolata | Ciliophora | Cyclotrichium_like_organism | - |
| Eukaryota | TSAR | Alveolata | Ciliophora | Heterotrichea | Yes |
| Eukaryota | TSAR | Alveolata | Ciliophora | Karyorelictea | - |
| Eukaryota | TSAR | Alveolata | Ciliophora | Litostomatea | Yes |
| Eukaryota | TSAR | Alveolata | Ciliophora | Nassophorea | Yes |
| Eukaryota | TSAR | Alveolata | Ciliophora | Oligohymenophorea | Yes |
| Eukaryota | TSAR | Alveolata | Ciliophora | Phyllopharyngea | Yes |
| Eukaryota | TSAR | Alveolata | Ciliophora | Plagiopylea | Yes |
| Eukaryota | TSAR | Alveolata | Ciliophora | Prostomatea_1 | Yes |
| Eukaryota | TSAR | Alveolata | Ciliophora | Prostomatea_2 | Yes |
| Eukaryota | TSAR | Alveolata | Ciliophora | Prostomatea_3 | Yes |
| Eukaryota | TSAR | Alveolata | Ciliophora | Spirotrichea | Yes |

| Domain | Supergroup | Division | Subdivision | Class | Sequenced by PacBio |
|---|---|---|---|---|---|
| Eukaryota | TSAR | Alveolata | Colponemidia | Colponemidia_X | - |
| Eukaryota | TSAR | Alveolata | Dinoflagellata | Dinophyceae | Yes |
| Eukaryota | TSAR | Alveolata | Dinoflagellata | Dinophyta_X | Yes |
| Eukaryota | TSAR | Alveolata | Dinoflagellata | Noctilucophyceae | - |
| Eukaryota | TSAR | Alveolata | Dinoflagellata | Oxyrrhea | - |
| Eukaryota | TSAR | Alveolata | Dinoflagellata | Syndiniales | Yes |
| Eukaryota | TSAR | Alveolata | Perkinsea | Perkinsea_X | - |
| Eukaryota | TSAR | Alveolata | Perkinsea | Perkinsida | Yes |
| Eukaryota | TSAR | Rhizaria | Cercozoa | Cercozoa_X | - |
| Eukaryota | TSAR | Rhizaria | Cercozoa | Chlorarachniophyceae | Yes |
| Eukaryota | TSAR | Rhizaria | Cercozoa | Endomyxa-Ascetosporea | Yes |
| Eukaryota | TSAR | Rhizaria | Cercozoa | Endomyxa | Yes |
| Eukaryota | TSAR | Rhizaria | Cercozoa | Endomyxa-Gromiidea | Yes |
| Eukaryota | TSAR | Rhizaria | Cercozoa | Endomyxa-Phytomyxea | Yes |
| Eukaryota | TSAR | Rhizaria | Cercozoa | Endomyxa | Yes |
| Eukaryota | TSAR | Rhizaria | Cercozoa | Filosa | Yes |
| Eukaryota | TSAR | Rhizaria | Cercozoa | Filosa-Granofilosea | Yes |
| Eukaryota | TSAR | Rhizaria | Cercozoa | Filosa-Imbricatea | Yes |
| Eukaryota | TSAR | Rhizaria | Cercozoa | Filosa-Metromonadea | - |
| Eukaryota | TSAR | Rhizaria | Cercozoa | Filosa-Sarcomonadea | Yes |
| Eukaryota | TSAR | Rhizaria | Cercozoa | Filosa-Thecofilosea | Yes |
| Eukaryota | TSAR | Rhizaria | Cercozoa | Novel-clade-10-12 | Yes |
| Eukaryota | TSAR | Rhizaria | Cercozoa | Phaeodarea | Yes |
| Eukaryota | TSAR | Rhizaria | Foraminifera | Foraminifera_X | - |
| Eukaryota | TSAR | Rhizaria | Foraminifera | Globothalamea | Yes |
| Eukaryota | TSAR | Rhizaria | Foraminifera | Monothalamids | Yes |
| Eukaryota | TSAR | Rhizaria | Foraminifera | Tubothalamea | Yes |
| Eukaryota | TSAR | Rhizaria | Radiolaria | Acantharea | Yes |
| Eukaryota | TSAR | Rhizaria | Radiolaria | Polycystinea | Yes |
| Eukaryota | TSAR | Rhizaria | Radiolaria | RAD-A | Yes |
| Eukaryota | TSAR | Rhizaria | Radiolaria | RAD-B | Yes |
| Eukaryota | TSAR | Rhizaria | Radiolaria | RAD-C | Yes |
| Eukaryota | TSAR | Rhizaria | Radiolaria | Radiolaria_X | Yes |
| Eukaryota | TSAR | Stramenopiles | Bigyra | Bigyra_X | - |

| Domain | Supergroup | Division | Subdivision | Class | Sequenced by PacBio |
|---|---|---|---|---|---|
| Eukaryota | TSAR | Stramenopiles | Bigyra | Opalozoa | Yes |
| Eukaryota | TSAR | Stramenopiles | Bigyra | Sagenista | Yes |
| Eukaryota | TSAR | Stramenopiles | Gyrista | Aurearenophyceae | - |
| Eukaryota | TSAR | Stramenopiles | Gyrista | Bolidophyceae | Yes |
| Eukaryota | TSAR | Stramenopiles | Gyrista | Chrysista | Yes |
| Eukaryota | TSAR | Stramenopiles | Gyrista | Chrysomerophyceae | - |
| Eukaryota | TSAR | Stramenopiles | Gyrista | Chrysophyceae | Yes |
| Eukaryota | TSAR | Stramenopiles | Gyrista | Developea | - |
| Eukaryota | TSAR | Stramenopiles | Gyrista | Diatomeae | Yes |
| Eukaryota | TSAR | Stramenopiles | Gyrista | Dictyochophyceae | Yes |
| Eukaryota | TSAR | Stramenopiles | Gyrista | Eustigmatophyceae | Yes |
| Eukaryota | TSAR | Stramenopiles | Gyrista | Gyrista_X | Yes |
| Eukaryota | TSAR | Stramenopiles | Gyrista | Hyphochytriomyceta | Yes |
| Eukaryota | TSAR | Stramenopiles | Gyrista | MOCH-1 | - |
| Eukaryota | TSAR | Stramenopiles | Gyrista | MOCH-2 | Yes |
| Eukaryota | TSAR | Stramenopiles | Gyrista | MOCH-4 | Yes |
| Eukaryota | TSAR | Stramenopiles | Gyrista | Pelagophyceae | Yes |
| Eukaryota | TSAR | Stramenopiles | Gyrista | Peronosporomycetes | Yes |
| Eukaryota | TSAR | Stramenopiles | Gyrista | Phaeophyceae | Yes |
| Eukaryota | TSAR | Stramenopiles | Gyrista | Phaeothamniophyceae | - |
| Eukaryota | TSAR | Stramenopiles | Gyrista | Picophagea | - |
| Eukaryota | TSAR | Stramenopiles | Gyrista | Pinguiophyceae | - |
| Eukaryota | TSAR | Stramenopiles | Gyrista | Pirsoniales | Yes |
| Eukaryota | TSAR | Stramenopiles | Gyrista | Raphidophyceae | Yes |
| Eukaryota | TSAR | Stramenopiles | Gyrista | Schizocladiophyceae | - |
| Eukaryota | TSAR | Stramenopiles | Gyrista | Synchromophyceae | - |
| Eukaryota | TSAR | Stramenopiles | Gyrista | Xanthophyceae | Yes |
| Eukaryota | TSAR | Stramenopiles | Stramenopiles_X | Stramenopiles-Group-2 | - |
| Eukaryota | TSAR | Stramenopiles | Stramenopiles_X | Stramenopiles-Group-3 | Yes |
| Eukaryota | TSAR | Stramenopiles | Stramenopiles_X | Stramenopiles-Group-4 | - |
| Eukaryota | TSAR | Stramenopiles | Stramenopiles_X | Stramenopiles-Group-6 | - |
| Eukaryota | TSAR | Stramenopiles | Stramenopiles_X | Stramenopiles-Group-7 | - |
| Eukaryota | TSAR | Stramenopiles | Stramenopiles_X | Stramenopiles-Group-8 | - |
| Eukaryota | TSAR | Stramenopiles | Stramenopiles_X | Stramenopiles-Group-9 | Yes |

| Domain | Supergroup | Division | Subdivision | Class | Sequenced by PacBio |
|---|---|---|---|---|---|
| Eukaryota | TSAR | Stramenopiles | Stramenopiles_X | Stramenopiles_XX | Yes |
| Eukaryota | TSAR | Telonemia | Telonemia_X | Telonemia_XX | Yes |

**Supplementary Table 3.** Details of the short-read metabarcoding datasets used in this study. ID refers to the dataset ID in metaPR2[28]. Number of ASVs is obtained after processing the raw reads with DADA2[29].

| No. | ID | Dataset name | Ref | Env level 1 | Env level 2 | Description | Location | Sampling date | Sequencing technology | Fwd primer | Rev primer | Number of ASVs | Bioproject | Repository |
|---|---|---|---|---|---|---|---|---|---|---|---|---|---|---|
| 1 | 1 | Ocean Sampling Day | [30] | marine | euphotic | global coastal waters | global | 2014-06-21 | Illumina | TAReuk454FWD1 | V4 18S Next.Rev | 12451 | PRJEB8682 | ENA |
| 2 | 2 | Ocean Sampling Day | [30] | marine | euphotic | global coastal waters | global | 2015-06-21 | Illumina | TAReuk454FWD1 | V4 18S Next.Rev | 11083 | PRJEB8682 | ENA |
| 3 | 3 | Ocean Sampling Day | [30] | marine | euphotic | global coastal waters | global | 2014-06-21 | Illumina | TAReuk454FWD1 | V4 18S Next.Rev | 13405 | PRJEB8682 | ENA |
| 4 | 34 | Malaspina Expedition | [25] | marine | euphotic + aphotic | global marine waters | global | 2010-2011 | Illumina | TAReuk454FWD1 | TAReukREV3 | 17656 | PRJEB23771 | ENA |
| 5 | 35 | Malaspina Expedition | [31] | marine | euphotic | global marine waters | global | 2010-2011 | Illumina | TAReuk454FWD1 | TAReukREV3 | 15338 | PRJEB23913 | ENA |
| 6 | 53 | Biomarks | [32] | marine | euphotic | European coastal waters | coast of Europe | 2010-2011 | 454 | TAReuk454FWD1 | TAReukREV3 | 9097 | PRJEB9133 | NCBI |
| 7 | 69 | Mariana Trench 1 | [26] | marine | aphotic | deep sea trench | Mariana Trench | 2016 | Illumina | TAReuk454FWD1 | TAReukREV3 | 5842 | SRP141405 | NCBI |
| 8 | 70 | Mariana Trench 2 | [33] | marine | aphotic | deep sea trench | Mariana Trench | 2016 | Illumina | 3NDF | V4_euk_R2 | 1115 | PRJNA399026 | NCBI |
| 9 | 150 | Saint-Charles River | [34] | terrestrial | freshwater | river | Canada | 2016-2017 | Illumina | E572F | E1009R | 5584 | PRJEB36925 | NCBI |
| 10 | 183 | Lake Fuxian | [35] | terrestrial | freshwater | lake | China | 2015 | Illumina | NSF573 | NSR951 | 3224 | PRJNA534173 | NCBI |
| 11 | 185 | Lake Chaohu | [36] | terrestrial | freshwater | lake | China | 2014-2015 | Illumina | NSF573 | NSR951 | 1733 | PRJNA534176, PRJNA330896 | NCBI |
| 12 | 195 | Lake Baikal | [37] | terrestrial | freshwater | lake | Siberia | 2013 | Illumina | TAReuk454FWD1 | TAReukREV3 | 1125 | PRJEB24415 | ENA |
| 13 | 196 | Chevreuse | [38] | terrestrial | freshwater | ponds | France | 2012 | 454 | EK-565F | UNonMet | 807 | PRJNA259710 | NCBI |
| 14 | 197 | Mountain Lakes | [39] | terrestrial | freshwater | high-altitude lakes | Austria, Chile, Ethiopia | 2013 | Illumina | TAReuk454FWD1 | TAReukREV3 | 25935 | SRP065150 | NCBI |
| 15 | 198 | Lake Garda | [40] | terrestrial | freshwater | alpine lake | Italy | 2014-2015 | Illumina | TAReuk454FWD1 | V4 18S Next.Rev | 1173 | PRJEB36925 | ENA |
| 16 | 199 | Neotropical Soils | [41] | terrestrial | soil | Neotropical forest soils | Central and South America | 2012-2013 | Illumina | TAReuk454FWD1 | TAReukREV3 | 28474 | PRJNA317860 | NCBI |
| 17 | 200 | Parana River | [42] | terrestrial | freshwater | floodplain lakes | South America | 2012 | Illumina | Nex-18S-0587-F | Nex_18S_0964_R | 7540 | PRJEB23471; PRJEB23493 | ENA |
| 18 | 201 | Swiss Soils | [43] | terrestrial | soil | soil | Switzerland | 2013 | Illumina | TAReuk454FWD1 | TAReukREV3 | 49669 | PRJEB30010 | ENA |
| 19 | 203 | Scandinavian Lakes | [44] | terrestrial | freshwater | lakes | Scandinavia | 2011 | 454 | TAReuk454FWD1 | TAReukREV3 | 2782 | 7s6s8 | Dryad |
| 20 | 204 | Global Soils | [45] | terrestrial | soil | global soils | global | - | 454 | F515 | R119 | 663 | - | - |
| 21 | 205 | Tara Oceans V4 | [46] | marine | euphotic | global marine waters | global | 2009-2013 | Illumina | TAReuk454FWD1 | TAReukREV3 | 20180 | PRJEB6610 | ENA |
| 22 | 206 | Tara Arctic V4 | [46] | marine | euphotic | Arctic Ocean | Arctic | 2009-2013 | Illumina | TAReuk454FWD1 | TAReukREV3 | 2209 | PRJEB9737 | ENA |

**Supplementary Table 4.** Environmental diversity collected from public datasets. Reads were processed using DADA2[29]. OTUs were conservatively filtered to reduce the number of "contaminant" taxa being kept, and only OTUs that had at least 100 reads or were present in two or more distinct samples were retained.

| Sample type | No. of processed reads | No. of ASVs | No. of OTUs (97% similarity) | No. of OTUs post filteration |
|---|---|---|---|---|
| **soil** | 127,197,084 | 91,980 | 20,474 | 11,935 |
| **freshwater** | 11,985,510 | 16,396 | 10,067 | 3,788 |
| **marine euphotic** | 83,448,640 | 49,903 | 22,294 | 10,650 |
| **marine aphotic** | 12,534,646 | 78,806 | 4,664 | 2,890 |
| **Terrestrial (all)** | 139,182,594 | 108,376 | 30,541 | 15,723 |
| **Marine (all)** | 95,983,286 | 128,709 | 26,958 | 13,540 |
